# Targeting SerpinE1 reverses cellular features of Hutchinson-Gilford progeria syndrome

**DOI:** 10.1101/2021.11.05.467259

**Authors:** Giorgia Catarinella, Chiara Nicoletti, Andrea Bracaglia, Paola Procopio, Illari Salvatori, Marilena Taggi, Alberto Ferri, Cristiana Valle, Rita Canipari, Pier Lorenzo Puri, Lucia Latella

**Affiliations:** Epigenetics and Regenerative Medicine, IRCCS Fondazione Santa Lucia, Rome, Italy; Institute of Translational Pharmacology, National Research Council of Italy, Rome, Italy; Development, Aging and Regeneration Program, Sanford Burnham Prebys Medical Discovery Institute, La Jolla, CA, 92037, USA; DAHFMO, Unit of Histology and Medical Embryology, Sapienza, University of Rome, Rome, Italy; Department of Biology, University of Rome Tor Vergata, Rome, Italy; Department of Experimental Medicine, University of Rome “La Sapienza”, 00161 Rome, Italy

**Keywords:** HGPS, SerpinE1-PAI-1, TM5441, Aging

## Abstract

Hutchinson-Gilford progeria syndrome (HGPS) is a rare, fatal disease caused by Lamin A mutation, leading to altered nuclear architecture, loss of perinuclear heterochromatin and deregulated gene expression. HGPS patients eventually die by coronary artery disease and cardiovascular alterations. However, how deregulated transcriptional networks at the cellular level impact on the systemic disease phenotype is currently unclear. We have performed a longitudinal genome-wide analysis of gene expression in primary HGPS fibroblasts from patients at two sequential stages of disease that revealed a progressive activation of Rho signaling and SerpinE1, also known as Plasminogen Activator Inhibitor (PAI-1). siRNA-mediated downregulation or pharmacological inhibition of SerpinE1 by TM5441 could revert key pathological features of HGPS in patient-derived fibroblasts, including re-activation of cell cycle progression, reduced DNA damage signaling, decreased expression of pro-fibrotic genes and recovery of mitochondrial defects. These effects were accompanied by reduced levels of Progerin and correction of nuclear abnormalities. These data point to SerpinE1 as a novel potential effector of HGPS pathogenesis and target for therapeutic interventions.

## Introduction

Hutchinson-Gilford progeria syndrome (HGPS) is a lethal disease caused by the autosomal dominant *de novo* mutation (1824C>T, p.G608G) in exon 11 of the *Lamin A* (LMNA) gene that activates a splicing donor site resulting in the production of Progerin, a toxic LMNA variant (Eriksson et al. 2003; de Sandre-Giovannoli et al. 2003). Unlike LMNA, Progerin cannot be properly processed and remains permanently farnesylated, thereby exerting its toxic properties (Fong et al. 2004) and leading to features of premature aging (Dechat et al. 2007). HGPS patients share essential molecular and clinical features with physiological aging, making HGPS a condition that prematurely recapitulates the aging process (López-Otín et al. 2013; Gonzalo et al. 2017).

HGPS patients invariably die by an average of 14 years and no cure is currently available. Although genome-editing approaches have been recently published (Koblan et al. 2021; Santiago-Fernández et al. 2019; Beyret et al. 2019) their translation into clinic is not immediate. As such, the identification of pharmacological approaches with available compounds that could be immediately translated to patients is an urgent and imperative task, in order to counter disease progression, improve quality of life and extend lifespan of HGPS patients (Macicior et al. 2021; Guilbert et al. 2021).

Progerin accumulation provokes nuclear lamin thickening and increases cellular stiffness, eventually altering cellular and tissue functions (Dahl et al. 2006). At cellular level, HGPS patients display abnormal nuclear shape, loss of perinuclear heterochromatin, DNA damage accumulation, telomere shortening, genomic instability and acquisition of senescence phenotype (Benson et al. 2010; Goldman et al. 2004; Gonzalo & Kreienkamp 2015; Liu et al. 2006; Shumaker et al. 2006). The nuclear defects interfere with cellular functions including the inflammatory response, metabolic changes, proteostasis and mitochondrial dysfunction (López-Otín et al. 2013). Altogether, the cellular defects are responsible for the pathological features of progeroid syndrome. In particular, loss of perinuclear heterochromatin causes alterations in gene expression. Lamins are essential organizers of genome topology, allowing a spatial distribution of chromatin inside the nucleus (Zheng et al. 2018). In HGPS patients as well as during physiological aging, global changes in the nuclear architecture and the consequent loss of perinuclear heterochromatin lead to altered patterns of gene expression (Oberdoerffer & Sinclair 2007); however, how deregulated transcriptional networks at the cellular level impact on the systemic disease phenotype remains unclear. Indeed, at the clinical level,

HGPS individuals display atherosclerosis, osteoporosis, loss of subcutaneous fat, reduced bone density and cardiovascular alterations resulting in myocardial infarction and stroke, the main causes of their premature death within the second decade of age (Hamczyk et al. 2019; Merideth et al. 2008; di Pasquale & Condorelli 2019). In particular, as altered vascular homeostasis appears the major cause of cardiovascular alterations, most of the research has focused on dysfunctional activities of endothelial or other vessel-associated cells types. Still, nuclear defects are present in all cell types of HGPS individuals, thereby prompting an interest on the potential contribution of other cell types to HGPS-related cardiovascular pathogenesis. Among these cells, fibroblasts are major candidates as they regulate angiogenesis and vascular homeostasis directly, by releasing soluble vasoactive factors, or indirectly, by altering extracellular cellular matrix (ECM) composition. Consistently, the majority of cardiovascular diseases are associated with excessive ECM deposition of pro-fibrotic components and vascular stiffening, which ultimately determine pathological cardiac fibrosis (Mohindra et al. 2021). In this regard, studies on available HGPS-derived primary fibroblasts provide an opportunity to discover potential interplay between fibroblast-derived alteration of ECM and endothelial/vascular dysfunction and to possibly identify targets for interventions that prevent cardiovascular diseases associated to HPGS. To this purpose, we have performed a longitudinal gene expression RNA-seq analysis from primary fibroblasts derived from HGPS patients at two sequential stages of disease progression, in order to capture transcriptional abnormalities that could potentially lead to pathogenic ECM alterations.

We identified altered Rho-SerpinE1 pathway in HGPS fibroblasts during disease progression and show that SerpinE1 is as key molecular player that is deregulated in HGPS cells and could provide a target for therapeutic interventions.

## Results

### Transcriptional signature during HGPS progression

To identify altered patterns of gene expression in HGPS fibroblasts during disease progression, we performed RNA-sequencing (RNA-seq) analysis in HGPS dermal fibroblasts isolated from HGPS patient at different stages of the disease, namely 2 year- and 8-year-old patients (2YO HGPS and 8YO HGPS, respectively). As a control, we used human fibroblasts from aged-matched healthy donors (2YO Ctrl and 8YO Ctrl). This approach provides a longitudinal gene expression RNA-seq analysis from primary cells of HGPS patients.

HGPS fibroblasts showed typical alterations in cell proliferation, leading to cell cycle arrest, as evaluated by decreased expression of *CyclinA2* (Fig. S1A), increased expression of *p16/INK4* (Fig. S1B) and *p21/Waf1* transcripts (Fig. S1C), as well protein levels (Fig. S1D, E). HGPS fibroblasts also showed flat morphology and exhibited spontaneous beta-Galactosidase activity (Fig. S1F, G), two typical features of cellular senescence. Moreover, HGPS fibroblasts displayed nuclear abnormalities and spontaneous activation of DNA damage signaling, as shown by detection of nuclear foci enriched with H2AX Phosphorylated in Serine 139 (γ-H2AX), NbsI Phosphorylated in Serine 343 (p-NbsI) (Fig. S1H-M) and 53BP1 (Fig. S1N, O) by immunofluorescence. Activation of DNA damage response was also detected by Western blot, revealing increased levels of γ-H2AX, p-NbsI and the downstream DNA damage effector, Serine15-phosphorylated 53 protein (Fig. S1P, Q). These data are consistent with the acquisition of senescence-associated features in HGPS fibroblasts.

RNA-seq analysis identified 1,744 and 2,059 differentially expressed genes (DEGs) from the 2YO HGPS vs. 2YO Ctrl, and 8YO HGPS vs. 8YO Ctrl comparisons, respectively. The majority of DEGs were shared between 2-year and 8-year-old HGPS samples, with 783 common genes between the two conditions (Fig. 1A). Heatmap showed that control samples (2YO Ctrl and 8YO Ctrl) clustered separately from HGPS samples (2YO HGPS and 8YO HGPS) (Fig. 1B). Gene ontology (GO) for the common DEGs revealed that the enriched categories were related to fibrosis, DNA damage, checkpoint activation and deregulation of the cell cycle, according to the progeroid phenotype of HGPS cells (Fig. 1C). Heatmap of top DEGs among Ctrl and HGPS could discriminate two discrete clusters of gene expression, which included both up-regulated and down-regulated genes and that were progressively modulated from 2 to 8-years-old HGPS cells (Fig. 1D). GO analysis on up-regulated genes revealed that the Rho signaling was significantly activated during HGPS progression, as compared to control cells (Fig. 1E). Conversely,

**Figure 1.**
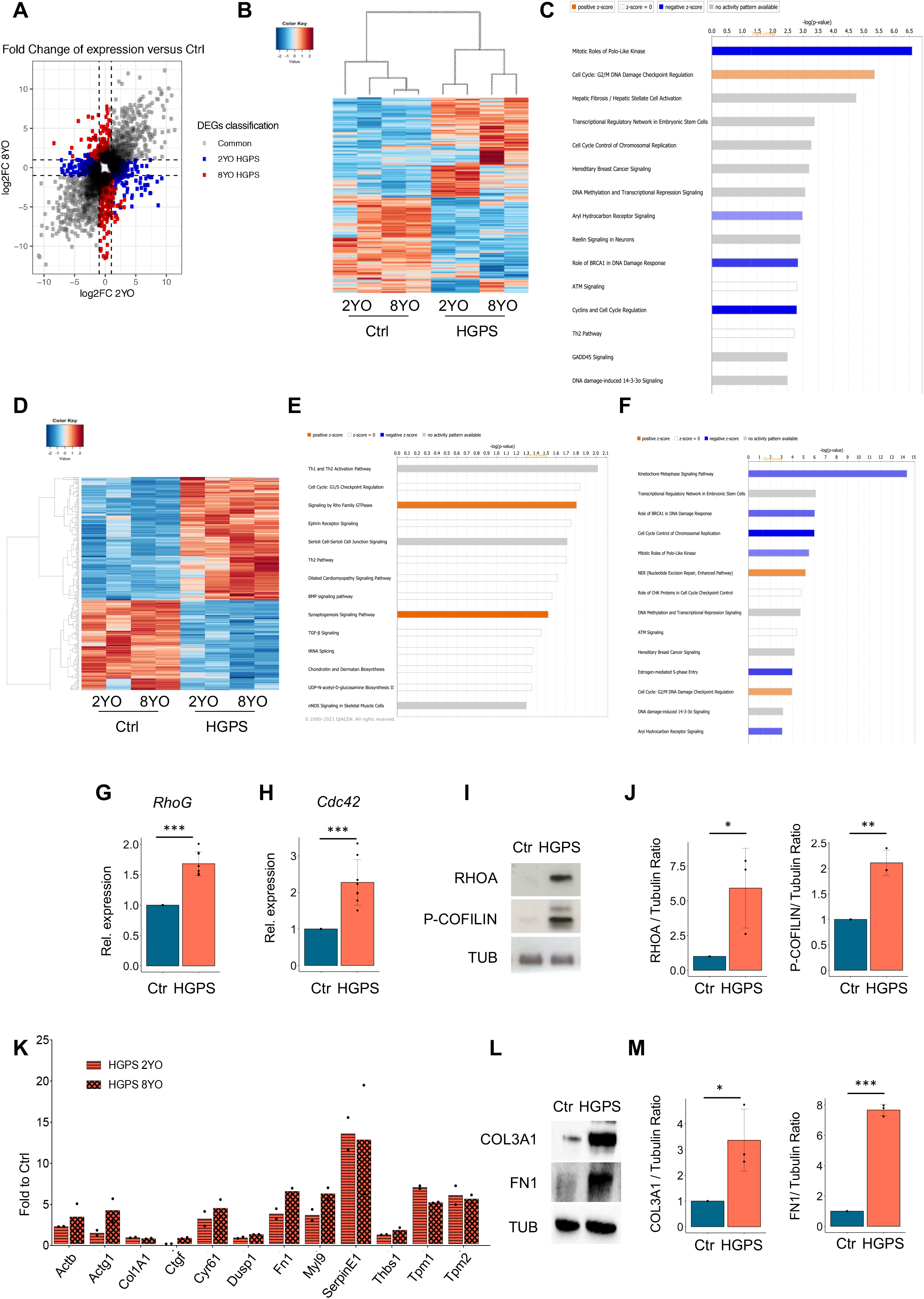
Transcriptional signature during HGPS progression. **A**. Scatterplot of differentially expressed genes (DEGs) of 2YO and 8YO HGPS fibroblasts. Common genes of 2YO and 8YO HGPS are shown in grey, 2YO HGPS unique genes in blue and 8YO HGPS unique genes in red. **B**. Heat map of DEGs from 2YO and 8YO HGPS vs. age matched control cells (2YO Ctrl and 8YO Ctrl). **C**. Ingenuity Pathway Analysis (IPA) showing enriched categories of common DEGs in 2YO and 8YO HGPS fibroblasts. **D**. Heatmap of 2YO and 8YO HGPS common DEGs vs. age-matched control cells (2YO Ctrl and 8YO Ctrl). **E**. IPA of a subset of DEGs from Fig. 1D that show a progressively higher expression rate compared to the healthy controls during disease progression (2 and 8 YO HGPS). **F**. IPA of a subset of DEGs from Fig. 1D that show a progressively lower expression rate compared to the healthy controls during disease progression (2 and 8 YO HGPS). **G, H**. qRT-PCR for *RhoG (n=7) and Cdc42 (n=7)* in control (Ctrl) and 8YO HGPS cells (HGPS). **I**. Cropped immunoblot for RhoA (RHOA), P-COFILIN and Tubulin (TUB) in control (Ctrl) and 8YO HGPS cells (HGPS). **J**. Plots represent RhoA/TUB ratio, P-COFILIN/TUB ratio based on the average for each experimental point (n = 3). **K**. qRT-PCR to reveal the fold induction of genes associated with fibrosis, *Actb* (actin beta), *Actg1* (actin gamma 1), *Col1A1* (Collagen Type I Alpha 1 Chain), *Ctgf* (Connective tissue growth factor), *Dusp1* (Dual Specificity Phosphatase 1), *Fn1* (Fibronectin1), *Myl9* (Myosin Light Chain9), *SerpinE1, Thbs1* (Thrombospondin 1), *Tpm1* (Tropomyosin 1), *Tpm2* (Tropomyosin 2) in 2YO and 8YO HGPS cells compared to control cells. **L**. Cropped immunoblot for Collagen3A1 (COL3A1), FIBRONECTIN1 (FN1) and Tubulin (TUB) in control (Ctrl) and 8YO HGPS cells (HGPS). **M**. Plots represent COL3A1/TUB ratio, FN1/TUB ratio based on the average for each experimental point, (n = 3).

GO on genes that progressively decline during the disease progression showed downregulation of gene networks associated to DNA repair and cell cycle progression (Fig. 1F). The activation of the Rho pathway was validated by q-RT-PCR for *RhoG* and *Cdc42* (Fig. 1G, H) and by Western blot analysis for RhoA and phospho-Cofilin, which were found activated in HGPS samples, as compared to control cells (Fig. 1I, J). Interestingly, previous studies have shown that activation of Rho signaling is associated to pathological alterations in ECM and development of fibrosis in multiple organs and tissues (Kawanami et al. 2016; Knipe et al. 2015; Cavalera et al. 2014; Tsou et al. 2014; Hartmann et al. 2015). Moreover, dysregulated Rho signaling has been implicated in HGPS-associated phenotypes (Kang et al. 2017a; Park et al. 2018).

As fibrosis was detected as a modulated pathway (Fig. 1C) and cardiovascular fibrosis is a hallmark of physiological aging and progeroid syndromes, we examined a panel of fibrogenic genes both by q-RT-PCR and by Western blot. We measured a significative upregulation of all analyzed genes associated with collagen deposition and fibrosis (Fig. 1K-M). In particular, our attention was captured by a putative downstream effector of Rho pathway - SerpinE1 (also known as PAI) (Samarakoon et al. 2008; Bryan et al. 2008; Kong et al. 2020) *-* whose expression was detected to increase up to 20-fold in HGPS fibroblasts by RNA-seq.

Interestingly, analysis of available datasets indicates that SerpinE1 was not upregulated in other cell types, such as endothelial cells, vascular smooth muscle cells and aorta cells (Bersini et al. 2020; von Kleeck et al. 2021), suggesting a fibroblast-specific induction of SerpinE1 in HGPS patients. Indeed, analysis of publicly available datasets generated from HGPS fibroblasts revealed a loss of LMNA binding to SerpinE1 promoter (McCord et al. 2013), with a consensual reduction in repressive chromatin marks (H3K27me3) at SerpinE1 promoter (Sebestyén et al. 2020). These data suggest that SerpinE1 expression is induced as a direct consequence of typical loss of heterochromatin by alteration of nuclear lamina, rather than secondary events developed in cultured HGPS fibroblasts.

As deregulated SerpinE1 activity has been largely associated to endothelial disfunction and cardiovascular diseases (Morrow et al. 2021; Jung et al. 2018), we decided to further explore the functional relationship between SerpinE1 up-regulation and pathological features in HGPS fibroblasts.

### SerpinE1 activity increases in HGPS primary cells

In order to investigate the aberrant activation of SerpinE1 in progeria samples, we first assessed its expression at transcriptional level. SerpinE1 expression increased in all HGPS primary cells that we analyzed compared to control cells (Fig. S2A). Moreover, functional activation of SerpinE1 was evaluated by reverse-zymography (Fig. S2B-E). In this assay, the presence of the inhibitory effect of unbound SerpinE1 was detected by the increased intensity of bands representing areas resistant to lysis (Fig. S2B). These data are in agreement with the higher levels of mRNA for SerpinE1 found in HGPS cells (Fig. S2A). Next, we evaluated the presence of Plasminogen Activator (PA) proteolytic activity by both zymography (Fig. S2C) and by chromogenic substrate assay (Fig. S2D). Furthermore, the “lytic areas” were plasminogen-dependent and were inhibited by amiloride, a specific inhibitor of uPA (Fig. S2E). Collectively, these data documented a decreased uPA activity in HGPS cells, as a consequence of increased expression of SerpinE1.

### SerpinE1 downregulation resumes proliferation and reduces DNA damage in HGPS

To investigate the function of SerpinE1 in HGPS fibroblasts, we evaluated the effect of SerpinE1 downregulation by siRNA (Fig. 2A). SerpinE1 downregulation could resume the cell cycle of HGPS fibroblasts, as revealed by EdU incorporation (Fig. 2B, C), which was associated with a reduction of DNA damage and Progerin levels (Fig. 2D, E) – two cardinal pathological features of HGPS (Sun et al. 2019).

**Figure 2.**
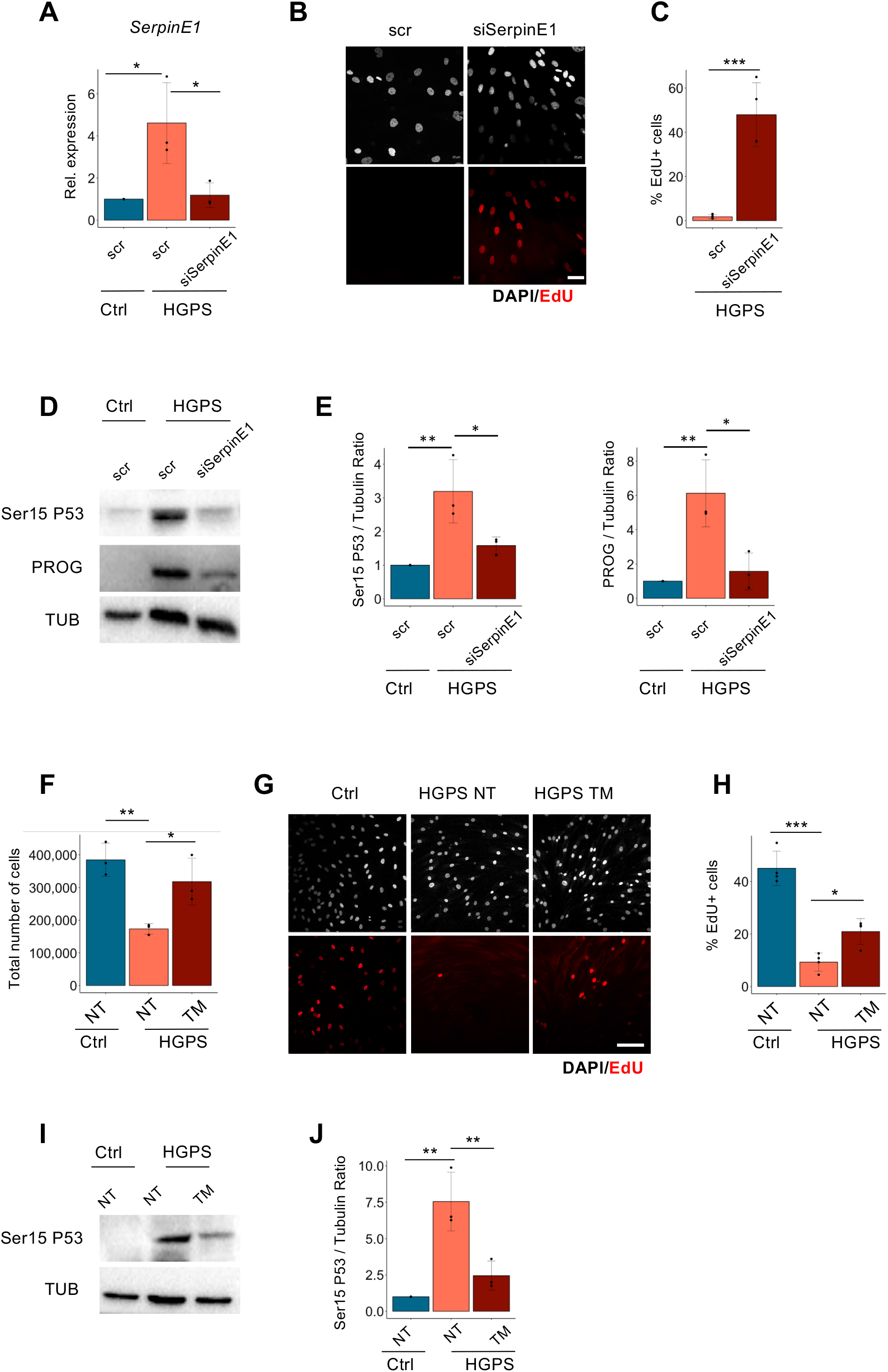
SerpinE1/PAI-1 downregulation resumes proliferation and reduces DNA damage in HGPS. **A**. qRT-PCR for *SerpinE1* transcript after 48h upon transfection in control cells (Ctrl) and HGPS transfected with scramble (scr) or siRNA for SerpinE1 (siSerpinE1), (n=3). **B**. Representative images relative to 8YO HGPS fibroblasts (HGPS) transfected with scramble (scr) or siRNA for SerpinE1 (siSerpinE1) analyzed five days post transfection and exposed to a 16 hours EdU pulse (red). Scale bar 50 μm. **C**. Quantification of the percentage EdU positive nuclei in 8YO HGPS fibroblasts (HGPS) transfected with scramble (scr) or siRNA for SerpinE1(siSerpinE1) analyzed five days post transfection and exposed to a 16 hours EdU (n=3). **D**. Cropped immunoblot for p53 phosphorylated at Serine 15 (Ser15 P53), Progerin (PROG) and Tubulin (TUB) as loading control in 8YO HGPS fibroblasts (HGPS) transfected with scramble (scr) or siRNA for SerpinE1 (siSerpinE1) analyzed five days post transfection. **E**. Plots represent Ser15 P53/TUB ratio and PROG/TUB ratio based on the average for each experimental point, (n = 3). **F**. Plot representing the total cell number of 8YO HGPS fibroblasts (HGPS) untreated and treated with TM5441 (TM) for 10 days compared to the age-matched control (Ctrl). **G**. Representative images relative to 8YO HGPS fibroblasts (HGPS) untreated and treated with TM5441 (TM) for 10 days and age-matched control (Ctrl) exposed to a 16 hours EdU pulse (red). Scale bar 50 μm. **H**. Quantification of the percentage EdU positive nuclei in 8YO HGPS fibroblasts (HGPS) untreated and treated with TM5441 (TM) for 10 days and age-matched control (Ctrl) exposed to a 16 hours EdU pulse, (n=4). **I**. Cropped immunoblot for p53 phosphorylated at Serine 15 (Ser15 P53) and Tubulin (TUB) as loading control in 8YO HGPS (HGPS) untreated and treated with TM5441(TM) and age matched control (Ctrl) cells. **J**. Plot represents Ser15 P53/TUB ratio based on the average for each experimental point (n = 3).

### TM5441 administration reverts HGPS pathological features

We next exploited the effect of pharmacological inhibition of SerpinE1 by an orally bioavailable compound, TM5441 (Piao et al. 2016). We initially verified the effect of TM5441 treatment, at the dosage of 10μM for 10 days, as already described (Sun et al. 2019), on SerpinE1 activity. Exposure to TM5441 increased uPA activity (Fig. S3A, B), leading to a partial recovery of activated metalloproteinase-2 (MMP2) in HGPS primary fibroblasts, as compared to untreated cells (Fig. S3C, D), demonstrating the efficacy of TM5441 treatment in reducing SerpinE1 activity in HGPS cells.

The effect of pharmacological inhibition of SerpinE1 activity by TM5441 was then tested on several pathological features exhibited by HGPS fibroblasts. We monitored the efficacy of TM5441 treatment in restoring proliferation in HGPS human fibroblasts. 8 YO HGPS human fibroblasts were untreated or treated with TM5441 then assessed for growth rate (Fig. 2F) and for their ability to incorporate 5-ethynyl-2**′**-deoxyuridine (EdU) (Fig. 2G, H). Thus, our data showed that SerpinE1 inhibition by TM5441 treatment could restore the proliferative potential of HGPS human fibroblasts. In addition, we observed a reduced DNA damage signaling in TM-treated HGPS cells as evaluated by reduced amount of Ser15-P53 levels (Fig 2I, J).

We further investigated the effects of TM5441 administration on additional features of HGPS, such as senescence, DNA damage accumulation and fibrosis. We cultured HGPS fibroblasts in the absence or in the presence of TM5441, administered for 10 days at the concentration of 10μM. Since HGPS cells acquire a precocious senescent phenotype, we monitored the ability of TM5441 in decelerating or mitigating the senescence mark revealed by Beta-Gal staining. We showed that SerpinE1 inhibition by TM5441 reduces the number of Beta-Gal-positive cells (Fig 3A, B) as well as lowers P16 levels (Fig 3C, D). Likewise, TM5441 treatment attenuated the activation of DNA damage response in HGPS fibroblasts, as measured by immunofluorescence for γ-H2AX (Fig 3E, F) and 53BP1 (Fig 3G, H). As γ-H2AX is only a proxy for DNA damage, we used alkaline Comet assay to monitor the DNA damage repair ability at the single-cell level, and showed that TM5441 treatment leads to a significant reduction of DNA lesions in HGPS fibroblasts (Fig. 3I).

**Figure 3.**
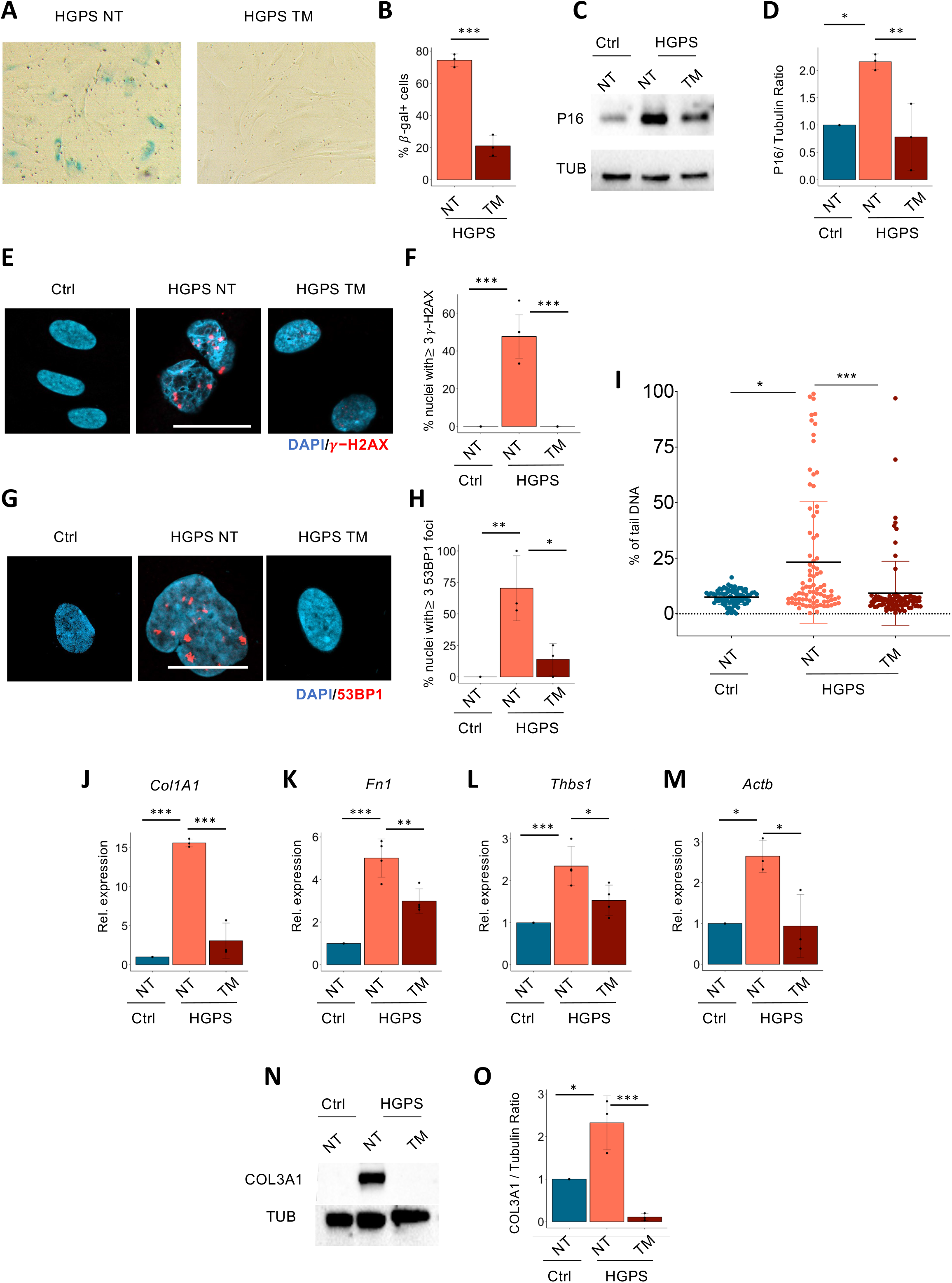
TM5441 treatment reverts pathological features associated to HGPS. **A**. Beta-Gal staining in 8YO HGPS fibroblasts untreated (HGPS NT) or treated with TM5441 (HGPS TM). **B**. Quantification of the percentage of beta-gal positive cells from 8YO HGPS fibroblasts untreated (HGPS NT) or treated with TM5441 (HGPS TM), (n=3). **C**. Cropped immunoblot for P16 and Tubulin (TUB) in 8YO HGPS (HGPS) untreated and treated with TM5441(TM) and age matched control (Ctrl) cells. **D**. Plot represents P16/TUB ratio based on the average for each experimental point, (n = 3). **E**. Representative images of immunofluorescence for γ-H2AX (red) and DAPI (blue) in 8YO HGPS fibroblasts untreated (HGPS NT) or treated with TM5441 (HGPS TM) and age-matched control (Ctrl). Scale bar 30 *μ*M. **F**. Quantification of the percentage of nuclei from 8YO HGPS fibroblasts untreated (HGPS NT) or treated with TM5441 (HGPS TM) and age-matched control (Ctrl) with at least three γ-H2AX foci, (n=3). **G**. Representative images of immunofluorescence for 53BP1 (red) and DAPI (blue) in 8YO HGPS fibroblasts untreated (HGPS NT) or treated with TM5441 (HGPS TM) and age-matched control (Ctrl). Scale bar 30 *μ*M. **H**. Quantification of the percentage of nuclei from 8YO HGPS fibroblasts untreated (HGPS NT) or treated with TM5441 (HGPS TM) and age-matched control (Ctrl) with at least three 53BP1 foci, (n=3). **I**. Alkaline comet assay performed in control fibroblasts (Ctrl), 8YO HGPS (HGPS) untreated and treated with TM5441 (TM) for 10 days. The percentage of tail DNA values is represented in the graphs. **J, K, L, M**. qRT-PCR for *Col1A1 (n=3), Fn1 (n=4), Thbs1 (n=4) and Actb (n=3)* in control (Ctrl) and 8YO HGPS (HGPS) untreated and treated with TM5441 (TM)). **N**. Cropped immunoblot for Collagen3A1 (COL3A1) and Tubulin (TUB) in 8YO HGPS (HGPS) untreated and treated with TM5441(TM) and age matched control (Ctrl) cells. **O**. Plot represents COL3A1/TUB ratio based on the average for each experimental point, (n = 3).

We next sought to investigate the effect of TM5441 on ECM dysregulation, which plays a significant role in HGPS progression (Harten et al. 2011). We showed that the administration of TM5441 leads to the reduction of ECM-related markers expression in HGPS cells, including a significant reduction in the levels of MMP-2 (Fig. S3C, D) and downregulation of *collagen1A1* (Fig. 3J), *fibronectin1* (Fig. 3K), *thrombospondin* 1 (Fig. 3L), *actin beta* (Fig. 3M) and Collagen3A1 (Fig 3N, O).

It was recently demonstrated that endothelial dysfunction caused by Progerin accumulation directly contributes to fibrosis and cardiac impairment in HGPS (Osmanagic-Myers et al. 2019). We then measured the capacity of TM5441 treatment to affect the delay in the acquisition of the progeria-associated phenotype such as nuclear blebbing. Normal (not blebbed) vs. abnormal (blebbed) nuclei were quantified, showing that the majority of HGPS cells treated with SerpinE1 inhibitor displayed a normal nuclear shape (Fig 4A, B). This evidence indicates that TM5441 administration is able to revert the morphological abnormalities associated to progeroid syndrome.

**Figure 4.**
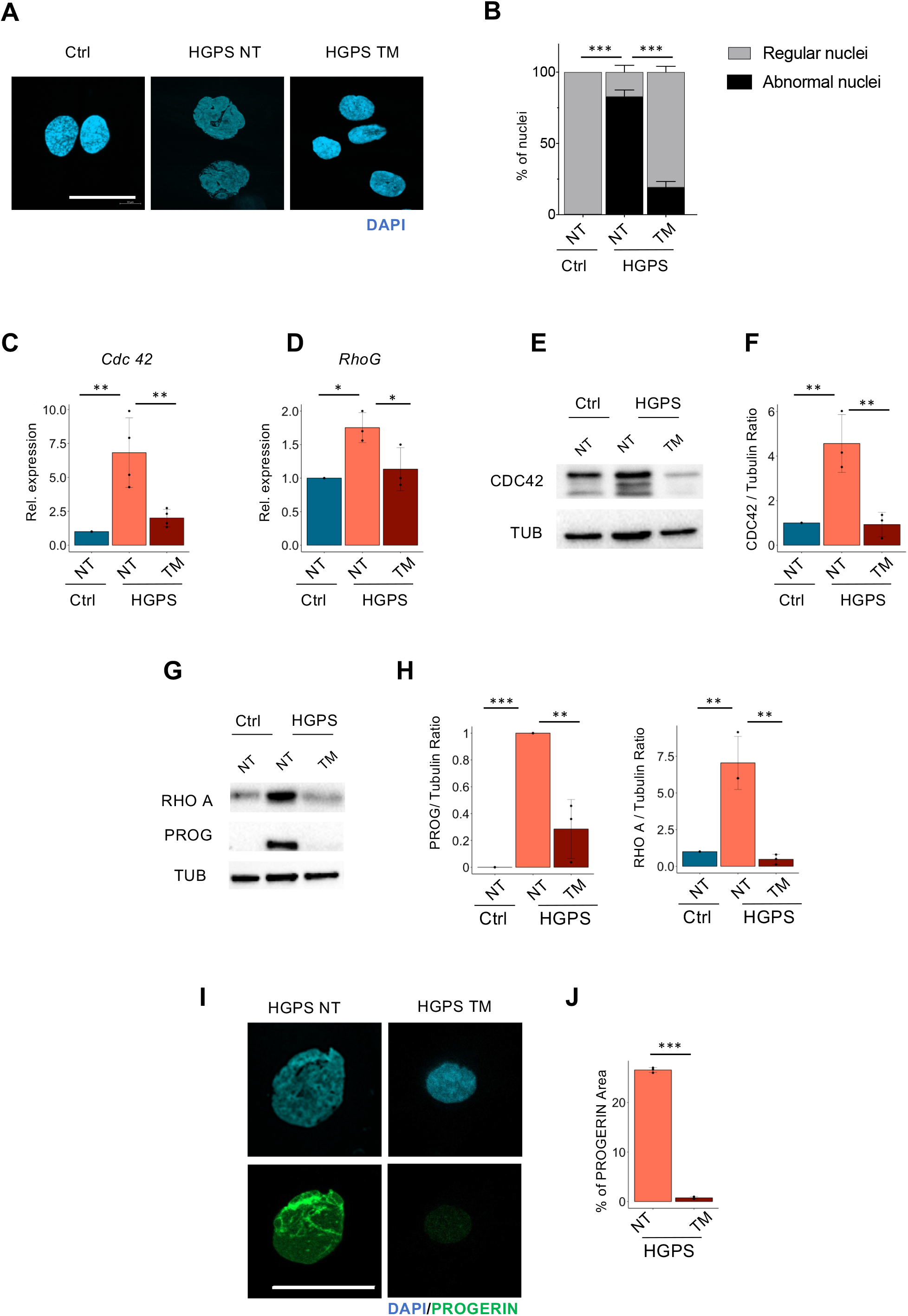
TM5441 treatment restores normal nuclear shape, the Rho pathway and prevents Progerin accumulation. **A**. Representative images of immunofluorescence for DAPI (blue) in 8YO HGPS fibroblasts untreated (HGPS NT) or treated with TM5441 (HGPS TM) and age-matched control (Ctrl). Scale bar 30 *μ*M. **B**. Quantification of regular and abnormal nuclei in control (Ctrl) and 8YO HGPS cells untreated (NT) or treated with TM5441 (TM), (n=3). **C, D**. qRT-PCR for *Cdc42 (n=4)* and *RhoG (n=3)* in control (Ctrl) and 8YO HGPS (HGPS) untreated and treated with TM5441 (TM). **E**. Cropped immunoblot for CDC42 and Tubulin (TUB) in 8YO HGPS (HGPS) untreated and treated with TM5441(TM) and age matched control (Ctrl) cells, (n=3). **F**. Plot represents *CDC42/*TUB ratio based on the average for each experimental point (n = 3). **G**. Cropped immunoblot for RHOA, Progerin (PROG) and Tubulin (TUB) in 8YO HGPS (HGPS) untreated and treated with TM5441(TM) and age matched control (Ctrl) cells. **H**. Plots represent PROG/TUB ratio and RHOA/TUB ratio based on the average for each experimental point (n = 3). **I**. Representative images of immunofluorescence for PROGERIN (green) and DAPI (blue) in 8YO HGPS fibroblasts untreated (HGPS NT) or treated with TM5441 (HGPS TM) and age-matched control (Ctrl). Scale bar 30 *μ*M. **J**. Quantification of PROGERIN area in control (Ctrl) and 8YO HGPS cells untreated (NT) or treated with TM5441 (TM), (n=3).

Nuclear lamin faces both intranuclear chromosomal organization and extranuclear cytoskeletal structure influencing migration polarization and cell mechano-properties (Lee et al. 2007; Lee et al. 2021). We therefore assessed whether TM5441 treatment might influence the activation of members of the Rho family of small GTPase that plays a key role in regulating cell polarization and migration. We measured a significant reduction of *Cdc42* (Fig 4C) and *RhoG* (Fig 4D) expression, as well as reduced amount of CDC42 (Fig 4E, F) and RHOA (Fig 4G, H) at protein level. Strikingly, treatment with TM5441 was also effective in reducing the accumulation of Progerin in HGPS-treated cells (Fig 4G-J).

### TM5441 treatment reverts mitochondrial defects in HGPS

HGPS are characterized by mitochondrial defects and mitochondrial dysfunction contributes to premature organ decline and aging in HGPS (Rivera-Torres et al. 2013). Yet, recovery of mitochondrial function has been proved to ameliorate HGPS phenotype (Kang et al. 2017b). We therefore, evaluated the ability of TM5441 in affecting mitochondrial functionality, by performing a Mito Stress Test using the Seahorse^®^XFe Technologies. This assay allows the analysis of a wide panel of mitochondrial bioenergetic parameters through the real-time measurement of oxygen consumption rate (OCR) in live cultured cells and enabled us to evaluate the differences in the energy profile between TM5441 treated and untreated HGPS cells. As described by representative OCR profiles, TM5441 treatment improved the bioenergetic performances of HGPS cells (Fig 5A). Data analysis showed a significant difference in basal respiration between treated and untreated HGPS cell, indeed, HPGS cells showed a compromission in basal OCR partially reverted by TM441 treatment (Fig 5B).

**Figure 5.**
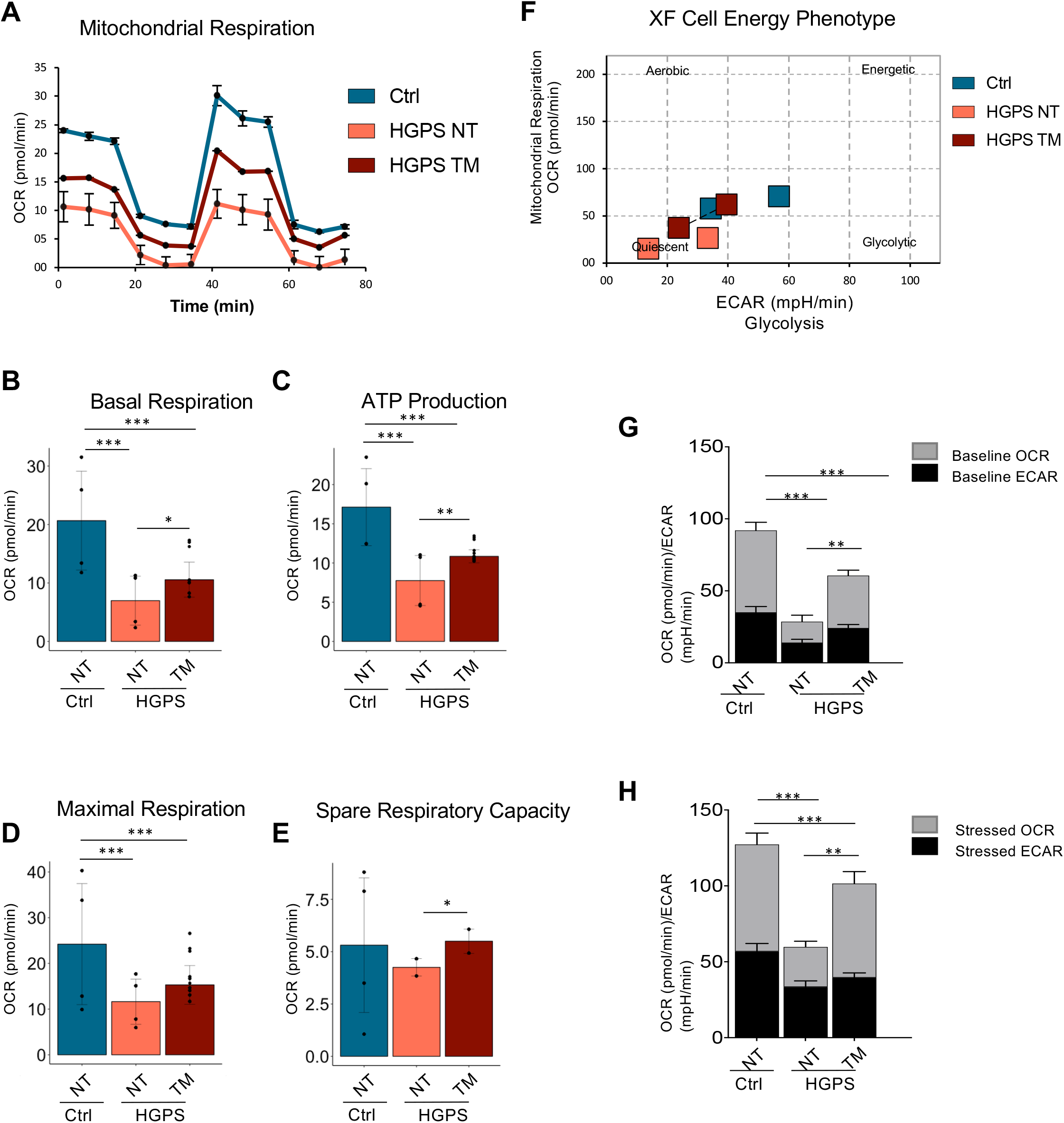
TM5441 treatment improves energy metabolism of HGPS. **A**. Representative data graphic of oxygen consumption rate (OCR) obtained from the Cell Mito Stress Test. Mitochondrial respiration expressed as normalized OCR of control, HGPS and HGPS TM5441-treated cells after the sequential addition of oligomycin, FCCP, rotenone and antimycin as indicated. **B**. The basal respiration which represents the oxygen consumption used to meet cellular ATP demand in resting condition. **C**. Mitochondrial ATP production, calculated as the OCR difference between the baseline and after the addition of oligomycin. **D**. The maximal respiration measured as the maximal oxygen consumption rate attained by adding the uncoupler FCCP. **E**. The spare respiratory capacity obtained from the analysis of from Maximal respiration. **F**. The cellular energy diagram showing the normalized oxygen consumption rate (OCR) and the normalized extracellular acidification rate (ECAR) of control, HGPS and HGPS TM5441-treated cells at baseline and stressed conditions. **G, H**. Metabolic potential of control, HGPS and HGPS TM5441-treated fibroblasts was calculated from unstressed (G) and stressed (H) OCR/ECAR over baseline.

We inferred the ATP production, by analyzing the difference between the basal oxygen consumption and values measured after oligomycin injection. HGPS fibroblasts showed a significant drop in ATP production that was partially reverted by TM5441 treatment (Fig. 5C). The addition of phenylhydrazone (FCCP), dissipating the protonic gradient, maximizes the electron flow through the mitochondrial transport chain and increases oxygen consumption which indicates the efficiency of the electron flow. The inability of HGPS cells to maximize the OCR following FCCP was partially abrogated by the TM441 treatment (Fig 5D). Similarly, the spare respiratory capacity (SRC), a measure of the ability of the cell to respond to increased energy demand was improved in HGPS cells, upon TM441 treatment (Fig. 5E). Indeed, FCCP mimics a physiological “energy demand” by stimulating the respiratory chain to operate at maximum capacity, which causes rapid oxidation of substrates to meet metabolic challenges.

These results are summarized by the Cell energy phenotype map (Fig. 5F) that clearly indicates a net improvement in the energy metabolism of HGPS cells treated with TM441 as further showed by the analysis of the OCR/ECAR (extracellular acidification rate) ratio performed with or without stressors (Fig. 5 G,H)

## Discussion

Our results revealed that Rho-SerpinE1 is aberrantly activated in HGPS fibroblasts and could be a potential target for pharmacological interventions in HGPS.

SerpinE1 functions as the primary inhibitor of plasminogen activators (PA) - tissue plasminogen activator (tPA) and urokinase (uPA) – thereby, in association with the broad family of matrix metalloproteinases (MMPs), affects fibrinolysis (Cesari et al. 2010; Ghosh & Vaughan 2012). Increased levels of SerpinE1 have been reported in metabolic associated diseases (Furukawa et al. 2004) as well as in increased occurrence of thrombosis (Meltzer et al. 2010) and development of atherosclerosis (Carratala et al. 2018), while lower SerpinE1 levels have been associated to protection against physiological aging (Eren et al. 2014; Khan et al. 2017). Intriguingly several reports show that SerpinE1 orchestrates the senescence-associated secretory phenotype (SASP) (Ozcan et al. 2016; Basisty et al. 2020) and regulates senescence and longevity in a mouse model of accelerated aging (Eren et al. 2014), pointing to SerpinE1 as potential player in cellular senescence (Kortlever et al. 2006). More recently, systemic SerpinE1 concentration was deemed a senescent-related biomarker that indicates an age-related process in non-human primates (Kavanagh et al. 2020). Hence, the SerpinE1 inhibitor TM5441 protects against cellular senescence (Ghosh et al. 2016), is effective against high-fat diet-induced obesity and adipocyte injury (Piao et al. 2016) and attenuates hypertension, cardiac hypertrophy, and periaortic fibrosis (Boe et al. 2013; Kaikita et al. 2001). These data indicate that TM5441 administration might provide a valid approach in preventing vascular aging and hypertension that are critical features of age-related pathologies (Bautista-Nino et al. 2016; Prakash et al. 2018).

We show that the treatment with SerpinE1 antagonist TM5441 ameliorates several pathological features in HGPS fibroblasts and could restore normal nuclear shape as well as prevent Progerin accumulation, presumably restoring the physiological activity of nuclear lamins.

As lamins regulate cellular mechano-sensing adaptive mechanisms, disruption of lamin activity and consequence nuclear abnormalities might directly cause DNA damage accumulation and altered gene expression (Cho et al. 2019), but can also promote activation of signaling implicated in mechano-transduction. Intriguingly, we also found that Rho signaling was activated in HGPS fibroblasts. The Rho-mediated signaling pathway influences a number of processes including motility, mitosis, and the entire cytoskeleton system that overall affect transcription regulation (Rajakyla & Vartiainen 2014). Remarkably, it has been previously shown that Rho-associated protein kinase (ROCK) is involved in mitochondrial reactive oxygen species (ROS) generation. ROCK inhibition leads to an amelioration of the HGPS phenotype (Kang et al. 2017b), also effective for the treatment of other age-related diseases (Park et al. 2018).

Currently, the only FDA-approved drug for HGPS boys is the farnesyltransferase inhibitor (FTI) Lonafarib (Gordon et al. 2012). However, many therapeutic approaches are developing to treat patients with HGPS (Macicior et al. 2021). The therapeutic strategies include gene therapies, which exploit CRISPR/Cas9 (Beyret et al. 2019; Santiago-Fernández et al. 2019), *in vivo* adenine base editors (ABEs) (Koblan et al. 2021) and small molecules that display different biological features. Among these, are included prenylation and methylation inhibitors, LMNA binders and modulators of the downstream detrimental outcomes associated to Progerin accumulation (Aguado et al. 2019). From the mono-drug treatment with Lonafarib clinical trial, that lower mortality rate (3.7% vs. 33.3%) after 2.2 years follow up (Gordon et al. 2018), the field has moved forward to multi-drug approaches, including two prenylation inhibitors Pravastatin and Zoledronate (Gordon et al. 2016). Lonafarnib monotherapy displays cardiovascular benefit in HGPS patients, but triple therapy does not provide additional benefit, suggesting that it Lonafarnib is the main responsible for lifespan extension. This data set the urgency to investigate additional candidate drugs or strategies aimed at depleting Progerin from the nucleus. Recent evidence shows that the Rapamycin analog Everolimus helps in reducing the amount of Progerin rescuing senescence features as well as nuclear morphology (DuBose et al. 2018). A new compound has been identified, named Remodelin, which ameliorates the HGPS cellular aberrations (Larrieu et al. 2014), leading to decreased genomic instability, as well as to an improved age-related phenotype in an established mouse model of HGPS (Balmus et al. 2018). Notably, Remodelin, which targets and inhibits the N-acetyltransferase NAT10 resulting in the rescue of the abnormal nuclear shape through microtubule rearrangement, is a potential therapeutic target for HGPS.

The inhibition of SerpinE1 by the TM5441 molecule is therefore a potential therapeutic strategy for combinatorial HGPS treatments. By targeting SerpinE1 we would expect to reduce senescence and fibrosis affecting cardiovascular complications occurring in HGPS (Kaikita et al. 2001; Olive et al. 2010; Osmanagic-Myers et al. 2019).

### Experimental Procedures Cell culture and treatments

To study DNA damage accumulation, we used human dermal fibroblasts obtained by the Progeria Research Foundation. We employed human fibroblasts isolated from 2YO (HGADFN003) and 8YO (HGADFN167, AG11513) HGPS patients, diagnosed by the detection of the c.1824C>T (p.Gly608Gly) heterozygous LMNA pathogenic variant. Other cell line (HGADFN 188, HGADFN127, HGADFN367, HGADFN169) were used to validate data. As a control we used fibroblasts derived from aged-matched healthy donors (2YO Ctrl - AG07095 - and 8YO Ctrl - GM08398 -) provided by Coriell Cell Repositories. Cells were cultured in growth medium, Dulbecco’s modified Eagles medium (DMEM) (Life Technologies) supplemented with 15% fetal bovine serum (FBS) (Hyclone) and penicillin/streptomycin at 37°C. Fibroblasts were cultured on 20 and 56 cm^2^ plates and allowed to reach 90% confluence, then trypsinized and splitted 1:2. For siRNA transfection, HGPS cells were transfected with siRNA for SerpinE1 (Ambion, ID #s10015), using jetOPTIMUS® (Polyplus, 117-01) according to manufacturer instructions; after 4 hours the medium was replaced with growth medium. To inhibit SerpinE1/PAI-1, we treated fibroblasts with TM5441 (Tocris) at concentration of 10 µM for 10 days, refreshed every other day.

### Immunofluorescence

In order to follow DNA damage accumulation, we performed immunofluorescence assays in HGPS and age-matched control cells plated on coverslips and fixed with 4% paraformaldehyde or methanol/acetone for 10 min at room temperature (RT). Fixed cells were permeabilized with 0.25% Triton X-100 in PBS and nonspecific antibody-binding sites were blocked by incubation with 4% BSA in PBS for 1 hour at RT. The following primary antibodies were used: γ-H2AX (Millipore, 05-636), P-Nbs1 (Epitomics), 53BP1 (Novus Biologicals, NB100-304). Immunostaining with primary antibodies was performed overnight (O/N) at 4°C. Antibody binding was revealed using species-specific secondary antibodies coupled to Alexa Fluor 488 or 594 (Molecular Probes, Eugene, CA, USA). Progerin levels were detected by Alexis human antibody (13A4, ALX-804-662-R200). DNA synthesis was evaluated by EdU incorporation following the manufacturer**’**s instructions (Invitrogen). Nuclei were visualized by DAPI (4**′**,6**′**-diamino-2-phenylindole). Images were acquired using Zeiss LSM5 Pascal confocal microscope. Fields reported in the figures are representative of all examined fields. Images were assembled using ImageJ software (NIH, Bethesda, MD, USA) and Photoshop software (Adobe Systems Incorporated, San Jose, CA, USA).

### Protein extraction and Western blot

Proteins were extracted with RIPA buffer supplemented with PMSF and protease inhibitor cocktail as described in (Latella et al. 2017) and resolved in SDS polyacrilamide gels, then transferred to nitrocellulose membranes (Bio-Rad). Western blot was performed using antibodies against the following proteins: Progerin (AbCam, Ab66587), P16 (Santa Cruz Biotechnology, C-20), P21 (Santa Cruz Biotechnology, C-19), Ser15 P53 (Cell Signaling), pNBS1 (Epitomics), γ-H2AX (Millipore, 05-636), RhoA (Cell Signaling), P-Cophillin (Cell Signaling), Collagen 3A1 (Santa Cruz, sc-271249), Fibronectin1 (Genetex, GTX112794), CDC42 (Genetex, GTX134588), and tubulin (Neo Markers, Ab4) as protein normalization marker. HRP-conjugated secondary antibodies were revealed with the ECL chemiluminescent kit (Amersham) following the manufacturer’s instructions. The signal was detected with a ChemiDoc MP Imaging System (Bio-Rad).

### RNA extraction and RT–PCR

Total RNA was extracted using TRI Reagent (Sigma), according to the manufacturer’s instructions. RNA (0.5–1 µg) was retrotranscribed using the TaqMan reverse transcription kit (Applied Bio-systems). qRT-PCR was performed to analyze relative gene expression levels using SYBR Green master mix (Applied Biosystems). Relative gene expression was normalized for TBP expression value and calculated using the 2^ΔΔCT^ method. Primers sequences are listed in Table 1.

**Table 1.**
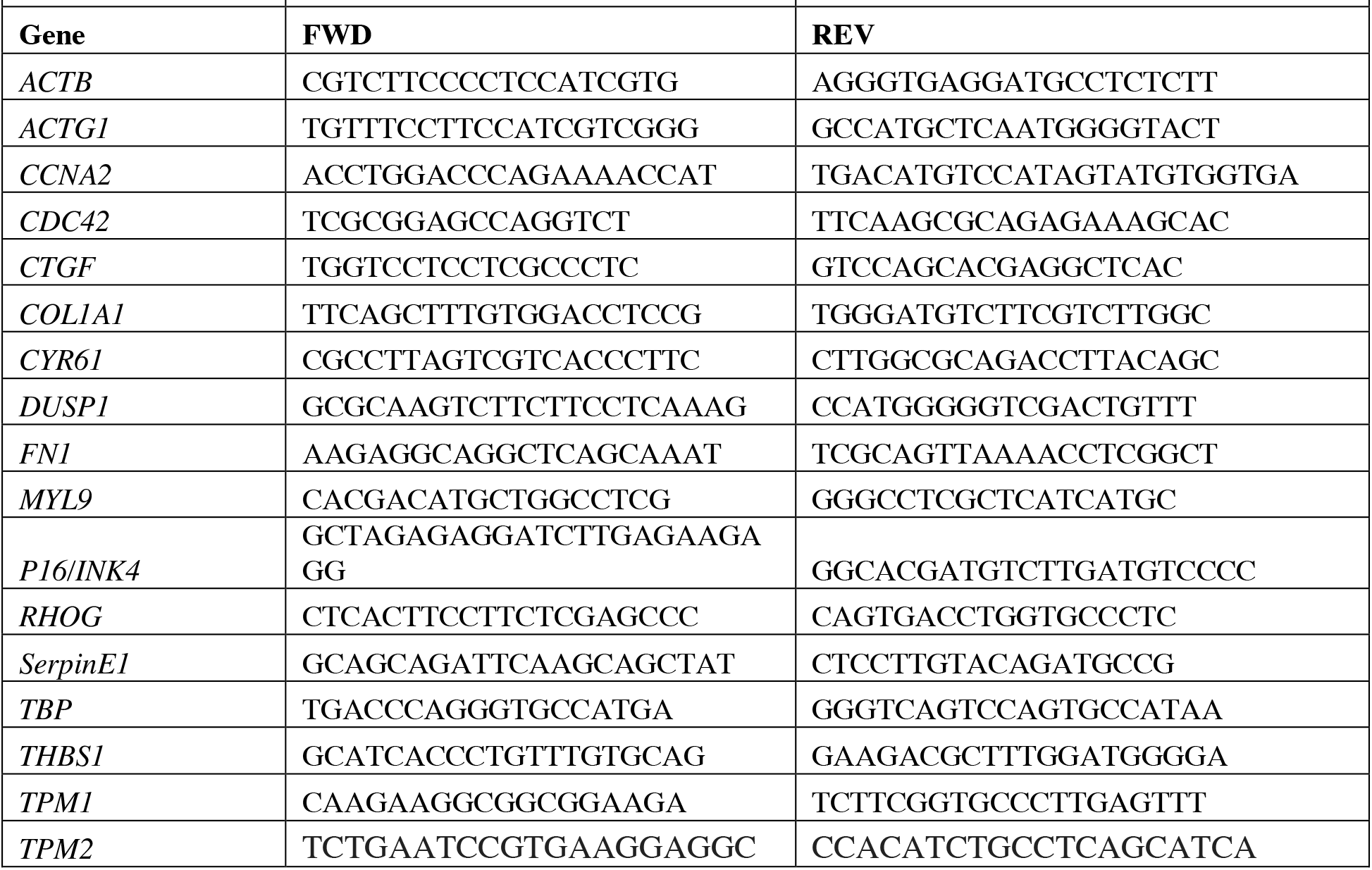
Primer list

### RNA-seq

RNA was isolated with Trizol from HGPS and age-matched control cells. One microgram of RNA was sent to the Institute of Applied Genomics (Udine, Italy) for deep sequencing. cDNA libraries were processed accordingly with the standard Illumina protocol and sequenced with the HiSeq2500 (4-plex run, 1×50 bp reads, about 30M reads/sample). Reads were aligned to the UCSC hg19 version of the human genome using Tophat2 ((Kim et al. 2013); v2.1.1), with parameters -g 1 --segment-length 24 -- library-type fr-secondstrand --no-coverage-search, and quantified to the hg19 UCSC genes with HTSeq-count ((Anders et al. 2015); v0.5.4p5) with parameters -m union -s yes.

Differential expression analysis was performed in R (v3.5.1) using DESeq2 ((Love et al. 2014); v1.20.0). Counts data from all conditions were filtered based on their raw count, keeping only those where the sum of the counts for all samples was higher than 1. Principal Component Analysis (PCA) was based on the 5% most variant genes between the different samples. Genes were considered differentially expressed with Benjamini–Hochberg adjusted p-value (FDR) <0.05. Ingenuity Pathway Analysis (IPA [Qiagen, http://www.qiagen.com/ ingenuity]) was used to perform gene ontology.

### Analysis of PA_s_ activities by zymography

Enzymatic activity of PA was assayed according to the method of Shimada and colleagues (Shimada et al. 1981) using a chromogenic substrate (substrate d-val-leulys-*p*-nitroanilide) assay. Samples were incubated with plasminogen, and the absorbance generated at 405 nm, related to PA activity, was normalized to cells present in the culture dish. To characterize the type of PA present in the samples, aliquots of conditioned medium were separated by 10% sodium dodecyl sulfate polyacrylamide gel electrophoresis (SDS-PAGE) under non-reducing conditions. After electrophoresis, gel was washed in 2,5% TritonX-100 for 30 minutes and then in water for 30 minutes. PA was then visualized by placing the Triton X-100 washed gel on a casein-agar-plasminogen underlay, as previously described (Catizone et al. 2010). All the bands were plasminogen dependent. PAI-1 was detected by reverse zymographic assays. 0.05 U/ml human urokinase (Serono, Denens, Switzerland) was added to the casein-agar-plasminogen underlay as previously described (Pepper et al. 1990). Molecular weights were calculated from the position of pre-stained molecular weight markers subjected to electrophoresis in parallel lanes.

### Gelatin zymography for matrix MMPs detection

Gelatinolytic activity of conditioned media was assayed as previously described (Catizone et al. 2010). Briefly, 20 μL aliquots of conditioned media were fractionated by 10% SDS-polyacrylamide gel electrophoresis in the presence of 0.1% gelatin under non-reducing conditions. Following electrophoresis, the gels were washed twice in 2.5% Triton X-100 for 30 min at room temperature to remove SDS. The gels were incubated at 37 °C overnight in substrate buffer, stained with 0.5% Coomassie Brilliant Blue R250 and distained in 30% methanol and 10% glacial acetic acid (vol/vol).

### Beta-galactosidase (beta-Gal) staining

HGPS fibroblasts were fixed in 4% formaldehyde and incubated at 37°C without CO_2_ with the following solution: 1mg/ml *X*-*gal* (5-bromo-4-chloro-3-indolyl-b-D-galactopyranoside), 5mM potassium ferrocyanide, 5mM potassium ferricyanide, 150 mM NaCl, 2 mM MgCl2 in 40mM citric acid/sodium phosphate pH 6.0. Staining was evident within 24 hours.

### Morphometric Analysis

To examine the overall percentage of blebbed nuclei in cells not treated and treated with TM5441, we performed a morphometric analysis in which nuclei were classified as blebbed if they contained three or more lobulations. Two independent observers performed this classification in a blind manner and the independent data sets were then averaged.

### Alkaline Comet Assay

2 YO and 8 YO HGPS were untreated or treated with TM5441 for 10 days. Single cells were analyzed for DNA breaks as previously described (Simonatto et al. 2013) employing the ‘TriTek Cometscore version 2.0’ software (Tritek Corporation, Sumerduck, VA, USA). The percentage of tail DNA was used as the measure of DNA damage. One hundred cells for each experimental point were scored.

### Bioenergetic analysis

Mitochondrial function and energy phenotype were determined using a Seahorse XF96e Analyzer (Seahorse Bioscience - Agilent, Santa Clara CA, USA). The fibroblasts were plated at the density of 15 × 10^4^. Mitochondrial stress test was performed according to Agilent’s recommendations. Briefly, growth medium was replaced with XF test medium (Eagle’s modified Dulbecco’s medium, 0 mM glucose, pH=7.4; Agilent Seahorse) supplemented with 1 mM pyruvate, 10 mM glucose and 2 mM L-glutamine. Before the assay the fibroblasts were incubated in 37° C incubator without CO_2_ for 1 h to allow to pre-equilibrate with the assay medium. The test was performed by measuring at first the baseline oxygen consumption rate (OCR), followed by sequential OCR measurements after injection of oligomycin (1,5 µM), carbonyl cyanide 4-(trifluoromethoxy) phenylhydrazone (1 µM) and Rotenone (0,5 µM) + Antimycin A (0,5 µM). This allowed to obtain the key parameters of the mitochondrial function including basal respiration, ATP-linked respiration, maximal respiration and spare respiratory capacity.

Cell energy phenotype test was performed according to Agilent’s recommendations. Briefly, growth medium was replaced with XF test medium (Eagle’s modified Dulbecco’s medium, 0 mM glucose, pH=7.4; Agilent Seahorse) supplemented as described above. The Cell Energy Phenotype Test Kit measures the basal OCR and then OCR after the injection of the stressor mix (1 μM of Oligomycin and 1 μM of FCCP). Oligomycin inhibits mitochondrial ATP production causing a compensatory increase in glycolysis rate measured as the Extracellular Acidification Rate (ECAR). FCCP depolarizes the mitochondrial membrane forcing the OCR to the maximum values. OCR/ECAR ratio, determined from the normalized OCR and normalized ECAR, indicating the cellular metabolic potential, were evaluated from the Cell Energy Phenotype Test data. All data were analyzed with XFe Wave software.

### Statistical analysis and reproducibility

Data are presented as mean ± SD of at least three independent biological replicates. The number of independent experimental replications are reported in the Figure legends (n).

Statistical analysis was conducted using Prism 7.0 A software (Pad Software). All data meet the assumptions of the tests (e.g., normal distribution). Statistical significance was determined using unpaired, two-tailed Student’s *t* test to compare the means of two groups, while One-way ANOVA or Two-way ANOVA were used, respectively, for comparison among the different groups and to examines the influence of two different independent variables on dependent variable. Tukey’s test was used for multiple comparison analysis.

Statistical significance was defined as P < 0.05 (*), P < 0.01 (**), and P < 0.001 (***). Immunofluorescence images are representative of at least 3 different experiments.

## Acknowledgements

We thank the Progeria Research Foundation (PRF) for providing primary human fibroblasts isolated from HGPS patients. We thank Emanuela Aleo at the Institute of Applied Genomics in Udine, Italy, for the RNA-seq library preparation and sequencing; Dr. A.Sacco and D. Palacios for critical reading of the manuscript and suggestions. This work was supported by Progetto Bandiera Epigenomica (EPIGEN) MIUR-CNR, R01 AR064873, AFM Research grant #20568, Italian Ministry of Health n. PE-2016-02363049 and H2020-MSCA-ITN-2019 grant # 860034 to LL.; C.N. is the recipient of the American Heart Association post-doctoral Fellowship 19POST34450187.

## Author Contributions

Project conception and design L.L.; Cell culture and treatments G.C., A.B., P.P.; Bioinformatics Analysis C.N.; Zymography M.T. and R.C.; Alkaline Comet assay A.B. Seahorse experiments G.C., I.S. and A.F. All authors discussed and interpreted data. Manuscript writing P.L.P. and L.L. Funding acquisition L.L.

## Supporting Informations

**Supplementary Figure 1.**
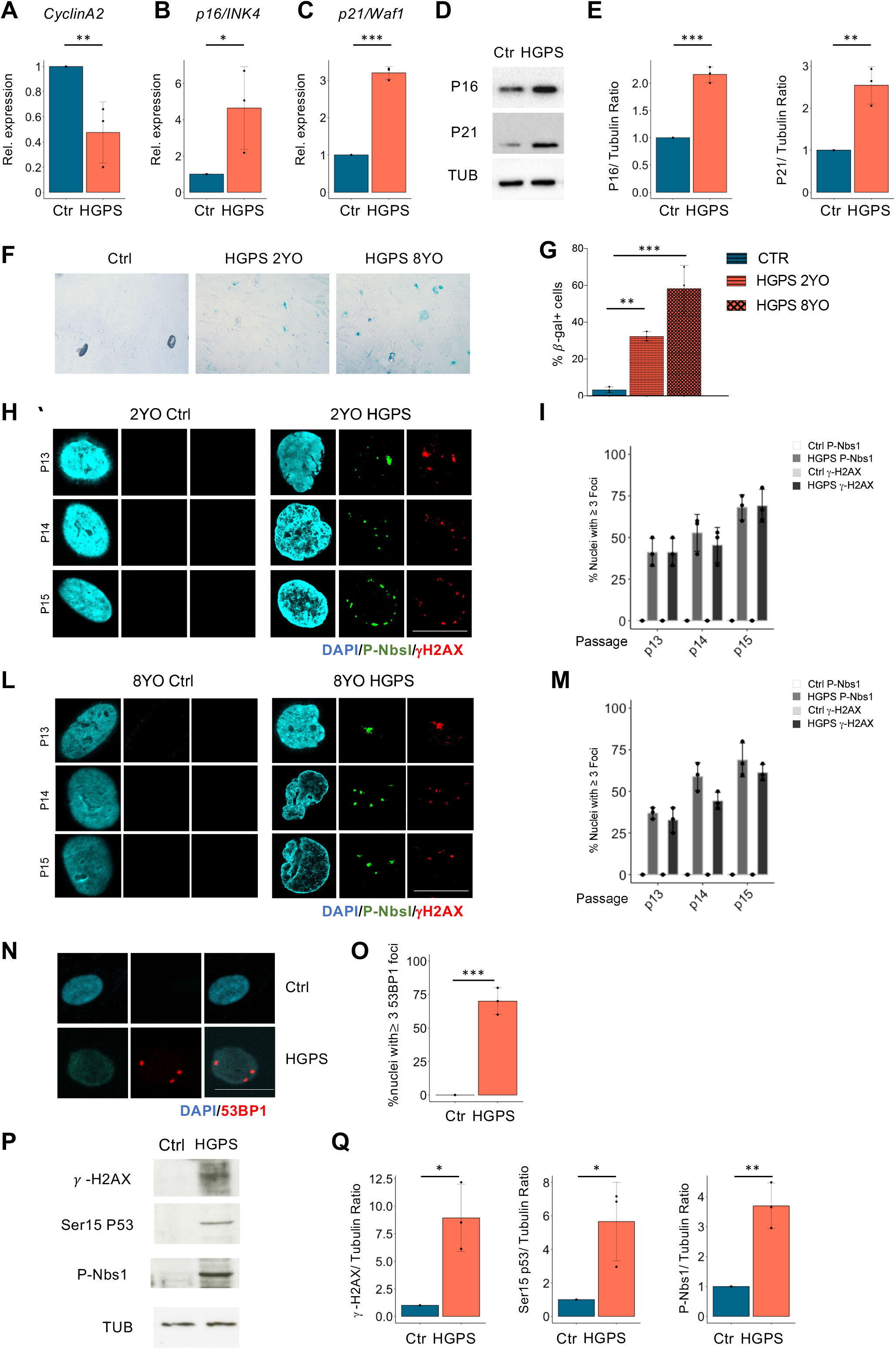
DNA damage accumulates precociously in cultured primary human HGPS compared to control fibroblasts. **A, B, C**. qRT-PCR for *CyclinA2* (n=3), *p16/INK4* (n=3) and *p21/Waf1 (n=3)* in control (Ctrl) and 8YO HGPS cells (HGPS). **D**. Cropped immunoblot for P16, P21 and Tubulin (TUB) in control (Ctrl) and 8YO HGPS cells (HGPS). **E**. Plots represent P16/TUB ratio, P21/TUB ratio based on the average for each experimental point, (n = 3). **F**. Beta-Gal staining in Ctrl, 2YO and 8YO HGPS fibroblasts. **G**. Quantification of the percentage of beta-gal positive cells of Ctrl, 2YO and 8YO HGPS fibroblasts **H**. Representative images of immunostaining for Phospho-NbsI (P-Nbs1) (green), γ-H2AX (red) and DAPI (blue) in 2YO HGPS and age matched control fibroblasts (2YO Ctrl) at different passage (p13, p14 and p15). Scale bar 50μM. **I**. Quantification of the percentage of nuclei with at least three foci positive for P-Nbs1 and γ-H2AX, (n=3). **L**. Representative images of immunostaining for P-NbsI (green), γ-H2AX (red) and DAPI (blue) in 8YO HGPS and age matched control fibroblasts (8YO Ctrl) at different passage (p13, p14 and p15). Scale bar 50μM. **M**. Quantification of the percentage of nuclei with at least three foci positive for P-NbsI and γ-H2AX, (n=3). **N**. Representative images of immunostaining for 53BP1 (red) and DAPI (blue) in HGPS fibroblasts isolated from 8YO patient (HGPS) and age matched control fibroblasts (Ctrl). Scale bar 30μM. **O**. Quantification of the percentage of nuclei with at least three foci positive for 53BP1, (n=3). **P**. Cropped immunoblot for γ-H2AX, p53 phosphorylated at Serine 15 (Ser15 P53), Phospho-NbsI (P-NBS1) and Tubulin (TUB) in 8YO HGPS (HGPS) and age matched control (Ctrl) cells. **Q**. Plots represent γ-H2AX /TUB ratio, Ser15 P53/TUB ratio and pNBS1/TUB ratio based on the average for each experimental point. (n = 3).

**Supplementary Figure 2.**
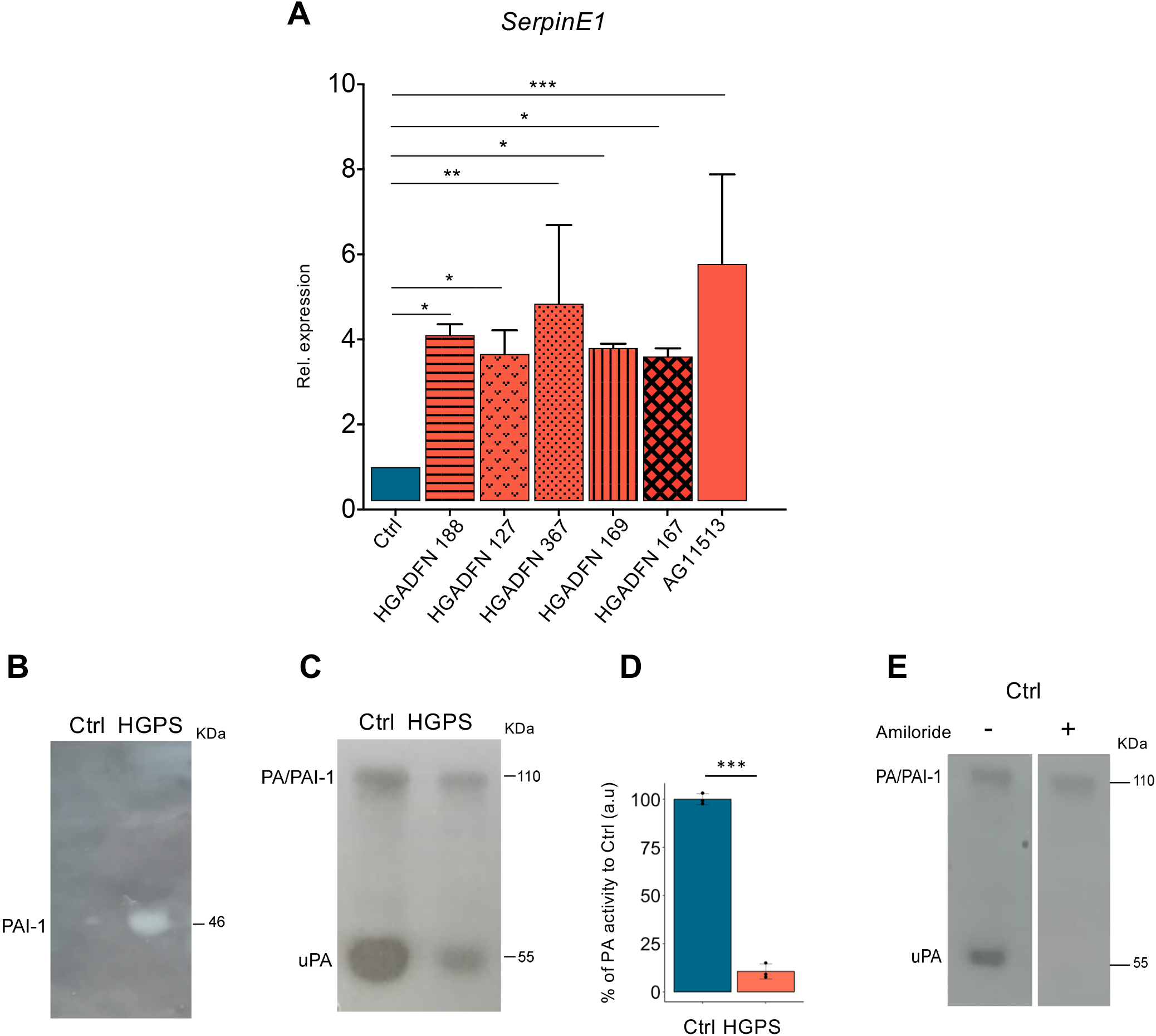
SerpinE1/PAI-1 expression and activity increase in HGPS primary cells. **A**. qRT-PCR for *SerpinE1* in control (Ctrl) and fibroblasts isolated from different HGPS patients: HGADFN 188, HGADFN127, HGADFN367, HGADFN169, HGADFN167 and AG11513, (n=3). **B**. Reverse zymography for SerpinE1 (PAI-1) in 8YO HGPS fibroblasts (HGPS) and age-matched control (Ctrl). **C**. Zymography for PA/PAI-1 complex and uPA in 8YO HGPS fibroblasts and age-matched control (Ctrl). **D**. Chromogenic substrate assay in 8YO HGPS fibroblasts (HGPS) and age-matched control (Ctrl). **E**. Amiloride treatment in control fibroblasts (Ctrl).

**Supplementary Figure 3.**
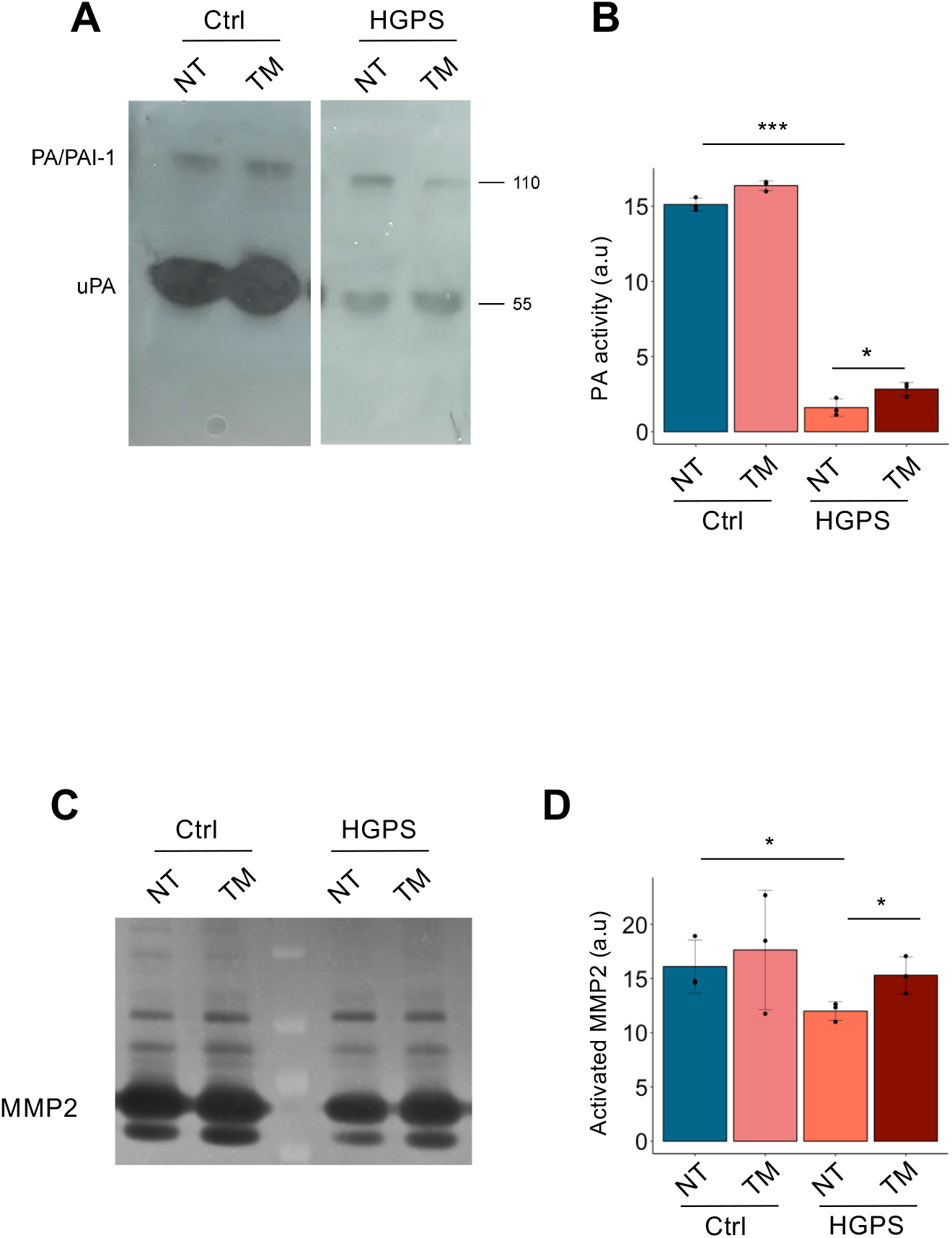
TM5441 is effective in restoring SerpinE1/PAI-1 activity in HGPS. **A**. Zymography for PA/PAI-1 complex and uPA in 8YO HGPS fibroblasts (HGPS) and age-matched control (Ctrl) untreated (NT) and treated with TM5441 (TM). **B**. Quantification of the PA activity, (n=3). **C**. Gelatin zymography for MMP2 detection in 8YO HGPS fibroblasts (HGPS) and age-matched control (Ctrl) untreated (NT) and treated with TM5441 (TM). **D**. Quantification of activated MMP2, (n=3).

## Data Availability Statement

The RNA-seq data supporting the findings of this study are openly available in the SRA database (https://www.ncbi.nlm.nih.gov/sra), with accession number PRJNA526216.

## References

Aguado J, Sola-Carvajal A, Cancila V, Revêchon G, Ong PF, Jones-Weinert CW, Wallén Arzt E, Lattanzi G, Dreesen O, Tripodo C, Rossiello F, Eriksson M & d’Adda di Fagagna F (2019) Inhibition of DNA damage response at telomeres improves the detrimental phenotypes of Hutchinson–Gilford Progeria Syndrome. Nature Communications 10. Available at: https://pubmed.ncbi.nlm.nih.gov/31740672/ [Accessed October 14, 2020].

Anders S, Pyl PT & Huber W (2015) HTSeq--a Python framework to work with high-throughput sequencing data. Bioinformatics 31, 166–169. Available at: https://www.ncbi.nlm.nih.gov/pubmed/25260700.

Balmus G, Larrieu D, Barros AC, Collins C, Abrudan M, Demir M, Geisler NJ, Lelliott CJ, White JK, Karp NA, Atkinson J, Kirton A, Jacobsen M, Clift D, Rodriguez R, Sanger Mouse Genetics P, Adams DJ & Jackson SP (2018) Targeting of NAT10 enhances healthspan in a mouse model of human accelerated aging syndrome. Nat Commun 9, 1700.

Basisty N, Kale A, Jeon OH, Kuehnemann C, Payne T, Rao C, Holtz A, Shah S, Sharma V, Ferrucci L, Campisi J & Schilling B (2020) A proteomic atlas of senescence-associated secretomes for aging biomarker development. PLoS Biol 18, e3000599. Available at: https://www.ncbi.nlm.nih.gov/pubmed/31945054.

Bautista-Nino PK, Portilla-Fernandez E, Vaughan DE, Danser AH & Roks AJ (2016) DNA Damage: A Main Determinant of Vascular Aging. Int J Mol Sci 17. Available at: https://www.ncbi.nlm.nih.gov/pubmed/27213333.

Benson EK, Lee SW & Aaronson SA (2010) Role of progerin-induced telomere dysfunction in HGPS premature cellular senescence. J Cell Sci 123, 2605–2612. Available at: https://www.ncbi.nlm.nih.gov/pubmed/20605919.

Bersini S, Schulte R, Huang L, Tsai H & Hetzer M (2020) Direct reprogramming of human smooth muscle and vascular endothelial cells reveals defects associated with aging and Hutchinson-Gilford progeria syndrome. eLife 9, 1–21. Available at: https://pubmed.ncbi.nlm.nih.gov/32896271/ [Accessed September 22, 2021].

Beyret E, Liao H, Yamamoto M, Hernandez-Benitez R, Fu Y, Erikson G, Reddy P & Izpisua Belmonte J (2019) Single-dose CRISPR-Cas9 therapy extends lifespan of mice with Hutchinson-Gilford progeria syndrome. Nature medicine 25, 419–422. Available at: https://pubmed.ncbi.nlm.nih.gov/30778240/ [Accessed July 27, 2021].

Boe AE, Eren M, Murphy SB, Kamide CE, Ichimura A, Terry D, McAnally D, Smith LH, Miyata T & Vaughan DE (2013) Plasminogen activator inhibitor-1 antagonist TM5441 attenuates Nomega-nitro-L-arginine methyl ester-induced hypertension and vascular senescence. Circulation 128, 2318–2324. Available at: https://www.ncbi.nlm.nih.gov/pubmed/24092817.

Bryan L, Paugh B, Kapitonov D, Wilczynska K, Alvarez S, Singh S, Milstien S, Spiegel S & Kordula T (2008) Sphingosine-1-phosphate and interleukin-1 independently regulate plasminogen activator inhibitor-1 and urokinase-type plasminogen activator receptor expression in glioblastoma cells: implications for invasiveness. Molecular cancer research : MCR 6, 1469–1477. Available at: https://pubmed.ncbi.nlm.nih.gov/18819934/ [Accessed September 22, 2021].

Carratala A, Martinez-Hervas S, Rodriguez-Borja E, Benito E, Real JT, Saez GT, Carmena R & Ascaso JF (2018) PAI-1 levels are related to insulin resistance and carotid atherosclerosis in subjects with familial combined hyperlipidemia. J Investig Med 66, 17–21. Available at: https://www.ncbi.nlm.nih.gov/pubmed/28822973.

Catizone A, Ricci G, Tufano MA, Perfetto B, Canipari R & Galdieri M (2010) Hepatocyte growth factor (HGF) modulates Leydig cell extracellular matrix components. J Androl 31, 306–313. Available at: https://www.ncbi.nlm.nih.gov/pubmed/19834131.

Cavalera M, Wang J & Frangogiannis N (2014) Obesity, metabolic dysfunction, and cardiac fibrosis: pathophysiological pathways, molecular mechanisms, and therapeutic opportunities. Translational research : the journal of laboratory and clinical medicine 164, 323–335. Available at: https://pubmed.ncbi.nlm.nih.gov/24880146/ [Accessed September 22, 2021].

Cesari M, Pahor M & Incalzi RA (2010) Plasminogen activator inhibitor-1 (PAI-1): a key factor linking fibrinolysis and age-related subclinical and clinical conditions. Cardiovasc Ther 28, e72–91. Available at: https://www.ncbi.nlm.nih.gov/pubmed/20626406.

Cho S, Vashisth M, Abbas A, Majkut S, Vogel K, Xia Y, Ivanovska IL, Irianto J, Tewari M, Zhu K, Tichy ED, Mourkioti F, Tang HY, Greenberg RA, Prosser BL & Discher DE (2019) Mechanosensing by the Lamina Protects against Nuclear Rupture, DNA Damage, and Cell-Cycle Arrest. Dev Cell 49, 920–935 e5. Available at: https://www.ncbi.nlm.nih.gov/pubmed/31105008.

Dahl KN, Scaffidi P, Islam MF, Yodh AG, Wilson KL & Misteli T (2006) Distinct structural and mechanical properties of the nuclear lamina in Hutchinson-Gilford progeria syndrome. Proc Natl Acad Sci U S A 103, 10271–10276. Available at: https://www.ncbi.nlm.nih.gov/pubmed/16801550.

Dechat T, Shimi T, Adam SA, Rusinol AE, Andres DA, Spielmann HP, Sinensky MS & Goldman RD (2007) Alterations in mitosis and cell cycle progression caused by a mutant lamin A known to accelerate human aging. Proc Natl Acad Sci U S A 104, 4955–4960. Available at: https://www.ncbi.nlm.nih.gov/pubmed/17360326.

DuBose AJ, Lichtenstein ST, Petrash NM, Erdos MR, Gordon LB & Collins FS (2018) Everolimus rescues multiple cellular defects in laminopathy-patient fibroblasts. Proc Natl Acad Sci U S A 115, 4206–4211.

Eren M, Boe AE, Murphy SB, Place AT, Nagpal V, Morales-Nebreda L, Urich D, Quaggin SE, Budinger GR, Mutlu GM, Miyata T & Vaughan DE (2014) PAI-1-regulated extracellular proteolysis governs senescence and survival in Klotho mice. Proc Natl Acad Sci U S A 111, 7090–7095. Available at: https://www.ncbi.nlm.nih.gov/pubmed/24778222.

Eriksson M, Brown WT, Gordon LB, Glynn MW, Singer J, Scott L, Erdos MR, Robbins CM, Moses TY, Berglund P, Dutra A, Pak E, Durkin S, Csoka AB, Boehnke M, Glover TW & Collins FS (2003) Recurrent de novo point mutations in lamin A cause Hutchinson-Gilford progeria syndrome. Nature 423, 293–298. Available at: https://www.ncbi.nlm.nih.gov/pubmed/12714972.

Fong LG, Ng JK, Meta M, Cote N, Yang SH, Stewart CL, Sullivan T, Burghardt A, Majumdar S, Reue K, Bergo MO & Young SG (2004) Heterozygosity for Lmna deficiency eliminates the progeria-like phenotypes in Zmpste24-deficient mice. Proc Natl Acad Sci U S A 101, 18111–18116. Available at: https://www.ncbi.nlm.nih.gov/pubmed/15608054.

Furukawa S, Fujita T, Shimabukuro M, Iwaki M, Yamada Y, Nakajima Y, Nakayama O, Makishima M, Matsuda M & Shimomura I (2004) Increased oxidative stress in obesity and its impact on metabolic syndrome. J Clin Invest 114, 1752–1761. Available at: https://www.ncbi.nlm.nih.gov/pubmed/15599400.

Ghosh AK, Rai R, Park KE, Eren M, Miyata T, Wilsbacher LD & Vaughan DE (2016) A small molecule inhibitor of PAI-1 protects against doxorubicin-induced cellular senescence. Oncotarget 7, 72443–72457. Available at: https://www.ncbi.nlm.nih.gov/pubmed/27736799.

Ghosh AK & Vaughan DE (2012) PAI-1 in tissue fibrosis. J Cell Physiol 227, 493–507. Available at: https://www.ncbi.nlm.nih.gov/pubmed/21465481.

Goldman RD, Shumaker DK, Erdos MR, Eriksson M, Goldman AE, Gordon LB, Gruenbaum Y, Khuon S, Mendez M, Varga R & Collins FS (2004) Accumulation of mutant lamin A causes progressive changes in nuclear architecture in Hutchinson-Gilford progeria syndrome. Proc Natl Acad Sci U S A 101, 8963–8968. Available at: https://www.ncbi.nlm.nih.gov/pubmed/15184648.

Gonzalo S & Kreienkamp R (2015) DNA repair defects and genome instability in Hutchinson-Gilford Progeria Syndrome. Current Opinion in Cell Biology 34, 75–83. Available at: https://pubmed.ncbi.nlm.nih.gov/26079711/ [Accessed October 14, 2020].

Gonzalo S, Kreienkamp R & Askjaer P (2017) Hutchinson-Gilford Progeria Syndrome: A premature aging disease caused by LMNA gene mutations. Ageing Res Rev 33, 18–29. Available at: https://www.ncbi.nlm.nih.gov/pubmed/27374873.

Gordon LB, Kleinman ME, Massaro J, D’Agostino Sr. RB, Shappell H, Gerhard-Herman M, Smoot LB, Gordon CM, Cleveland RH, Nazarian A, Snyder BD, Ullrich NJ, Silvera VM, Liang MG, Quinn N, Miller DT, Huh SY, Dowton AA, Littlefield K, Greer MM & Kieran MW (2016) Clinical Trial of the Protein Farnesylation Inhibitors Lonafarnib, Pravastatin, and Zoledronic Acid in Children With Hutchinson-Gilford Progeria Syndrome. Circulation 134, 114–125.

Gordon LB, Kleinman ME, Miller DT, Neuberg DS, Giobbie-Hurder A, Gerhard-Herman M, Smoot LB, Gordon CM, Cleveland R, Snyder BD, Fligor B, Bishop WR, Statkevich P, Regen A, Sonis A, Riley S, Ploski C, Correia A, Quinn N, Ullrich NJ, Nazarian A, Liang MG, Huh SY, Schwartzman A & Kieran MW (2012) Clinical trial of a farnesyltransferase inhibitor in children with Hutchinson-Gilford progeria syndrome. Proc Natl Acad Sci U S A 109, 16666–16671. Available at: https://www.ncbi.nlm.nih.gov/pubmed/23012407.

Gordon LB, Shappell H, Massaro J, D’Agostino Sr. RB, Brazier J, Campbell SE, Kleinman ME & Kieran MW (2018) Association of Lonafarnib Treatment vs No Treatment With Mortality Rate in Patients With Hutchinson-Gilford Progeria Syndrome. JAMA 319, 1687–1695.

Guilbert S, Cardoso D, Lévy N, Muchir A & Nissan X (2021) Hutchinson-Gilford progeria syndrome: Rejuvenating old drugs to fight accelerated ageing. Methods (San Diego, Calif.) 190, 3–12. Available at: https://pubmed.ncbi.nlm.nih.gov/32278808/ [Accessed September 22, 2021].

Hamczyk MR, Villa-Bellosta R, Quesada V, Gonzalo P, Vidak S, Nevado RM, Andres-Manzano MJ, Misteli T, Lopez-Otin C & Andres V (2019) Progerin accelerates atherosclerosis by inducing endoplasmic reticulum stress in vascular smooth muscle cells. EMBO Mol Med 11. Available at: https://www.ncbi.nlm.nih.gov/pubmed/30862662.

Harten IA, Zahr RS, Lemire JM, Machan JT, Moses MA, Doiron RJ, Curatolo AS, Rothman FG, Wight TN, Toole BP & Gordon LB (2011) Age-Dependent Loss of MMP-3 in Hutchinson– Gilford Progeria Syndrome. The Journals of Gerontology Series A: Biological Sciences and Medical Sciences 66A, 1201. Available at: /pmc/articles/PMC3193525/ [Accessed August 4, 2021].

Hartmann S, Ridley A & Lutz S (2015) The Function of Rho-Associated Kinases ROCK1 and ROCK2 in the Pathogenesis of Cardiovascular Disease. Frontiers in pharmacology 6. Available at: https://pubmed.ncbi.nlm.nih.gov/26635606/ [Accessed September 22, 2021].

Jung R, Simard T, Labinaz A, Ramirez F, di Santo P, Motazedian P, Rochman R, Gaudet C, Faraz M, Beanlands R & Hibbert B (2018) Role of plasminogen activator inhibitor-1 in coronary pathophysiology. Thrombosis research 164, 54–62. Available at: https://pubmed.ncbi.nlm.nih.gov/29494856/ [Accessed September 22, 2021].

Kaikita K, Fogo AB, Ma L, Schoenhard JA, Brown NJ & Vaughan DE (2001) Plasminogen activator inhibitor-1 deficiency prevents hypertension and vascular fibrosis in response to long-term nitric oxide synthase inhibition. Circulation 104, 839–844. Available at: https://www.ncbi.nlm.nih.gov/pubmed/11502712.

Kang HT, Park JT, Choi K, Choi HJ, Jung CW, Kim GR, Lee YS & Park SC (2017a) Chemical screening identifies ROCK as a target for recovering mitochondrial function in Hutchinson-Gilford progeria syndrome. Aging Cell. Available at: https://www.ncbi.nlm.nih.gov/pubmed/28317242.

Kang HT, Park JT, Choi K, Choi HJC, Jung CW, Kim GR, Lee YS & Park SC (2017b) Chemical screening identifies ROCK as a target for recovering mitochondrial function in Hutchinson-Gilford progeria syndrome. Aging Cell 16, 541–550. Available at: https://www.ncbi.nlm.nih.gov/pubmed/28317242.

Kavanagh K, Sherrill C, Ruggiero A, Block M, Vemuri R, Davis M & Olivier A (2020) Biomarkers of senescence in non-human primate adipose depots relate to aging. GeroScience, 1–10. Available at: https://link.springer.com/article/10.1007/s11357-020-00230-z [Accessed October 26, 2020].

Kawanami D, Matoba K & Utsunomiya K (2016) Signaling pathways in diabetic nephropathy. Histology and histopathology 31, 1059–1067. Available at: https://pubmed.ncbi.nlm.nih.gov/27094540/ [Accessed September 22, 2021].

Khan SS, Shah SJ, Klyachko E, Baldridge AS, Eren M, Place AT, Aviv A, Puterman E, Lloyd-Jones DM, Heiman M, Miyata T, Gupta S, Shapiro AD & Vaughan DE (2017) A null mutation in SERPINE1 protects against biological aging in humans. Sci Adv 3, eaao1617. Available at: https://www.ncbi.nlm.nih.gov/pubmed/29152572.

Kim D, Pertea G, Trapnell C, Pimentel H, Kelley R & Salzberg SL (2013) TopHat2: accurate alignment of transcriptomes in the presence of insertions, deletions and gene fusions. Genome Biol 14, R36. Available at: https://www.ncbi.nlm.nih.gov/pubmed/23618408.

von Kleeck R, Roberts E, Castagnino P, Bruun K, Brankovic S, Hawthorne E, Xu T, Tobias J & Assoian R (2021) Arterial stiffness and cardiac dysfunction in Hutchinson-Gilford Progeria Syndrome corrected by inhibition of lysyl oxidase. Life science alliance 4. Available at: https://pubmed.ncbi.nlm.nih.gov/33687998/ [Accessed September 22, 2021].

Knipe R, Tager A & Liao J (2015) The Rho kinases: critical mediators of multiple profibrotic processes and rational targets for new therapies for pulmonary fibrosis. Pharmacological reviews 67, 103–117. Available at: https://pubmed.ncbi.nlm.nih.gov/25395505/ [Accessed September 22, 2021].

Koblan L, Erdos M, Wilson C, Cabral W, Levy J, Xiong Z, Tavarez U, Davison L, Gete Y, Mao X, Newby G, Doherty S, Narisu N, Sheng Q, Krilow C, Lin C, Gordon L, Cao K, Collins F, Brown J & Liu D (2021) In vivo base editing rescues Hutchinson-Gilford progeria syndrome in mice. Nature 589, 608–614. Available at: https://pubmed.ncbi.nlm.nih.gov/33408413/ [Accessed July 27, 2021].

Kong HJ, Kwon EJ, Kwon OS, Lee H, Choi JY, Kim YJ, Kim W & Cha HJ (2020) Crosstalk between YAP and TGFβ regulates SERPINE1 expression in mesenchymal lung cancer cells. International Journal of Oncology 58, 111–121.

Kortlever RM, Higgins PJ & Bernards R (2006) Plasminogen activator inhibitor-1 is a critical downstream target of p53 in the induction of replicative senescence. Nat Cell Biol 8, 877–884. Available at: https://www.ncbi.nlm.nih.gov/pubmed/16862142.

Larrieu D, Britton S, Demir M, Rodriguez R & Jackson SP (2014) Chemical inhibition of NAT10 corrects defects of laminopathic cells. Science 344, 527–532.

Latella L, Dall’Agnese A, Boscolo FS, Nardoni C, Cosentino M, Lahm A, Sacco A & Puri PL (2017) DNA damage signaling mediates the functional antagonism between replicative senescence and terminal muscle differentiation. Genes Dev 31, 648–659. Available at: https://www.ncbi.nlm.nih.gov/pubmed/28446595.

Lee G, Han SB & Kim DH (2021) Cell-ECM contact-guided intracellular polarization is mediated via lamin A/C dependent nucleus-cytoskeletal connection. Biomaterials 268, 120548.

Lee JSH, Hale CM, Panorchan P, Khatau SB, George JP, Tseng Y, Stewart CL, Hodzic D & Wirtz D (2007) Nuclear Lamin A/C Deficiency Induces Defects in Cell Mechanics, Polarization, and Migration. Biophysical Journal 93, 2542–2552.

Liu Y, Rusinol A, Sinensky M, Wang Y & Zou Y (2006) DNA damage responses in progeroid syndromes arise from defective maturation of prelamin A. J Cell Sci 119, 4644–4649. Available at: https://www.ncbi.nlm.nih.gov/pubmed/17062639.

López-Otín C, Blasco MA, Partridge L, Serrano M & Kroemer G (2013) The Hallmarks of Aging. Cell 153, 1194–1217.

Love MI, Huber W & Anders S (2014) Moderated estimation of fold change and dispersion for RNA-seq data with DESeq2. Genome Biol 15, 550. Available at: https://www.ncbi.nlm.nih.gov/pubmed/25516281.

Macicior J, Marcos-Ramiro B & Ortega-Gutiérrez S (2021) Small-Molecule Therapeutic Perspectives for the Treatment of Progeria. International Journal of Molecular Sciences 22, 7190. Available at: https://www.mdpi.com/1422-0067/22/13/7190 [Accessed July 27, 2021].

McCord RP, Nazario-Toole A, Zhang H, Chines PS, Zhan Y, Erdos MR, Collins FS, Dekker J & Cao K (2013) Correlated alterations in genome organization, histone methylation, and DNA-lamin A/C interactions in Hutchinson-Gilford progeria syndrome. Genome Res 23, 260–269. Available at: https://www.ncbi.nlm.nih.gov/pubmed/23152449.

Meltzer ME, Lisman T, de Groot PG, Meijers JC, le Cessie S, Doggen CJ & Rosendaal FR (2010) Venous thrombosis risk associated with plasma hypofibrinolysis is explained by elevated plasma levels of TAFI and PAI-1. Blood 116, 113–121. Available at: https://www.ncbi.nlm.nih.gov/pubmed/20385790.

Merideth MA, Gordon LB, Clauss S, Sachdev V, Smith AC, Perry MB, Brewer CC, Zalewski C, Kim HJ, Solomon B, Brooks BP, Gerber LH, Turner ML, Domingo DL, Hart TC, Graf J, Reynolds JC, Gropman A, Yanovski JA, Gerhard-Herman M, Collins FS, Nabel EG, Cannon 3rd RO, Gahl WA & Introne WJ (2008) Phenotype and course of Hutchinson-Gilford progeria syndrome. N Engl J Med 358, 592–604. Available at: https://www.ncbi.nlm.nih.gov/pubmed/18256394.

Mohindra R, Agrawal D & Thankam F (2021) Altered Vascular Extracellular Matrix in the Pathogenesis of Atherosclerosis. Journal of cardiovascular translational research. Available at: https://pubmed.ncbi.nlm.nih.gov/33420681/ [Accessed July 27, 2021].

Morrow G, Whyte C & Mutch N (2021) A Serpin With a Finger in Many PAIs: PAI-1’s Central Function in Thromboinflammation and Cardiovascular Disease. Frontiers in cardiovascular medicine 8. Available at: https://pubmed.ncbi.nlm.nih.gov/33937363/ [Accessed September 22, 2021].

Oberdoerffer P & Sinclair DA (2007) The role of nuclear architecture in genomic instability and ageing. Nat Rev Mol Cell Biol 8, 692–702. Available at: https://www.ncbi.nlm.nih.gov/pubmed/17700626.

Olive M, Harten I, Mitchell R, Beers JK, Djabali K, Cao K, Erdos MR, Blair C, Funke B, Smoot L, Gerhard-Herman M, Machan JT, Kutys R, Virmani R, Collins FS, Wight TN, Nabel EG & Gordon LB (2010) Cardiovascular pathology in Hutchinson-Gilford progeria: correlation with the vascular pathology of aging. Arterioscler Thromb Vasc Biol 30, 2301–2309. Available at: https://www.ncbi.nlm.nih.gov/pubmed/20798379.

Osmanagic-Myers S, Kiss A, Manakanatas C, Hamza O, Sedlmayer F, Szabo PL, Fischer I, Fichtinger P, Podesser BK, Eriksson M & Foisner R (2019) Endothelial progerin expression causes cardiovascular pathology through an impaired mechanoresponse. J Clin Invest 129, 531–545. Available at: https://www.ncbi.nlm.nih.gov/pubmed/30422822.

Ozcan S, Alessio N, Acar MB, Mert E, Omerli F, Peluso G & Galderisi U (2016) Unbiased analysis of senescence associated secretory phenotype (SASP) to identify common components following different genotoxic stresses. Aging (Albany NY) 8, 1316–1329. Available at: https://www.ncbi.nlm.nih.gov/pubmed/27288264.

Park JT, Kang HT, Park CH, Lee YS, Cho KA & Park SC (2018) A crucial role of ROCK for alleviation of senescence-associated phenotype. Exp Gerontol 106, 8–15. Available at: https://www.ncbi.nlm.nih.gov/pubmed/29474864.

di Pasquale E & Condorelli G (2019) Endoplasmic reticulum stress at the crossroads of progeria and atherosclerosis. EMBO Mol Med 11. Available at: https://www.ncbi.nlm.nih.gov/pubmed/30902910.

Pepper MS, Belin D, Montesano R, Orci L & Vassalli JD (1990) Transforming growth factor-beta 1 modulates basic fibroblast growth factor-induced proteolytic and angiogenic properties of endothelial cells in vitro. J Cell Biol 111, 743–755. Available at: https://www.ncbi.nlm.nih.gov/pubmed/1696269.

Piao L, Jung I, Huh JY, Miyata T & Ha H (2016) A novel plasminogen activator inhibitor-1 inhibitor, TM5441, protects against high-fat diet-induced obesity and adipocyte injury in mice. Br J Pharmacol 173, 2622–2632. Available at: https://www.ncbi.nlm.nih.gov/pubmed/27339909.

Prakash A, Gordon LB, Kleinman ME, Gurary EB, Massaro J, D’Agostino Sr. R, Kieran MW, Gerhard-Herman M & Smoot L (2018) Cardiac Abnormalities in Patients With Hutchinson-Gilford Progeria Syndrome. JAMA Cardiol 3, 326–334. Available at: https://www.ncbi.nlm.nih.gov/pubmed/29466530.

Rajakyla EK & Vartiainen MK (2014) Rho, nuclear actin, and actin-binding proteins in the regulation of transcription and gene expression. Small GTPases 5, e27539. Available at: https://www.ncbi.nlm.nih.gov/pubmed/24603113.

Rivera-Torres J, Acín-Perez R, Cabezas-Sánchez P, Osorio FG, Gonzalez-Gómez C, Megias D, Cámara C, López-Otín C, Enríquez JA, Luque-García JL & Andrés V (2013) Identification of mitochondrial dysfunction in Hutchinson-Gilford progeria syndrome through use of stable isotope labeling with amino acids in cell culture. Journal of Proteomics 91, 466–477.

Samarakoon R, Higgins SP, Higgins CE & Higgins PJ (2008) TGF-β1-induced plasminogen activator inhibitor-1 expression in vascular smooth muscle cells requires pp60c-src/EGFRY845 and Rho/ROCK signaling. Journal of Molecular and Cellular Cardiology 44, 527–538.

de Sandre-Giovannoli A, Bernard R, Cau P, Navarro C, Amiel J, Boccaccio I, Lyonnet S, Stewart CL, Munnich A, le Merrer M & Levy N (2003) Lamin a truncation in Hutchinson-Gilford progeria. Science 300, 2055. Available at: https://www.ncbi.nlm.nih.gov/pubmed/12702809.

Santiago-Fernández O, Osorio FG, Quesada V, Rodríguez F, Basso S, Maeso D, Rolas L, Barkaway A, Nourshargh S, Folgueras AR, Freije JMP & López-Otín C (2019) Development of a CRISPR/Cas9-based therapy for Hutchinson–Gilford progeria syndrome. Nature Medicine 25, 423–426.

Sebestyén E, Marullo F, Lucini F, Petrini C, Bianchi A, Valsoni S, Olivieri I, Antonelli L, Gregoretti F, Oliva G, Ferrari F & Lanzuolo C (2020) SAMMY-seq reveals early alteration of heterochromatin and deregulation of bivalent genes in Hutchinson-Gilford Progeria Syndrome. Nature communications 11. Available at: https://pubmed.ncbi.nlm.nih.gov/33293552/ [Accessed September 22, 2021].

Shimada H, Mori T, Takada A, Takada Y, Noda Y, Takai I, Kohda H & Nishimura T (1981) Use of chromogenic substrate S-2251 for determination of plasminogen activator in rat ovaries. Thromb Haemost 46, 507–510. Available at: https://www.ncbi.nlm.nih.gov/pubmed/7197812.

Shumaker DK, Dechat T, Kohlmaier A, Adam SA, Bozovsky MR, Erdos MR, Eriksson M, Goldman AE, Khuon S, Collins FS, Jenuwein T & Goldman RD (2006) Mutant nuclear lamin A leads to progressive alterations of epigenetic control in premature aging, Available at: www.pnas.orgcgidoi10.1073pnas.0602569103.

Simonatto M, Marullo F, Chiacchiera F, Musaro A, Wang JY, Latella L & Puri PL (2013) DNA damage-activated ABL-MyoD signaling contributes to DNA repair in skeletal myoblasts. Cell Death Differ 20, 1664–1674. Available at: https://www.ncbi.nlm.nih.gov/pubmed/24056763.

Sun T, Ghosh AK, Eren M, Miyata T & Vaughan DE (2019) PAI-1 contributes to homocysteine-induced cellular senescence. Cell Signal 64, 109394. Available at: https://www.ncbi.nlm.nih.gov/pubmed/31472244.

Tsou P, Haak A, Khanna D & Neubig R (2014) Cellular mechanisms of tissue fibrosis. 8. Current and future drug targets in fibrosis: focus on Rho GTPase-regulated gene transcription. American journal of physiology. Cell physiology 307. Available at: https://pubmed.ncbi.nlm.nih.gov/24740541/ [Accessed September 22, 2021].

Zheng X, Hu J, Yue S, Kristiani L, Kim M, Sauria M, Taylor J, Kim Y & Zheng Y (2018) Lamins Organize the Global Three-Dimensional Genome from the Nuclear Periphery. Mol Cell 71, 802–815 e7. Available at: https://www.ncbi.nlm.nih.gov/pubmed/30201095.

